# Dynamic global acetylation remodeling during the yeast heat shock response

**DOI:** 10.1101/2025.01.10.632339

**Authors:** Rebecca E. Hardman-Kavanaugh, Aaron J. Storey, Tara N. Stuecker, Stephanie E. Hood, Gregory A. Barrett-Wilt, Venkata R. Krishnamurthi, Yong Wang, Stephanie D. Byrum, Samuel G. Mackintosh, Rick D. Edmondson, Wayne P. Wahls, Alan J. Tackett, Jeffrey A. Lewis

**Author notes:** Corresponding author: Jeffrey A. Lewis, Department of Biological Sciences, University of Arkansas, 850 W. Dickson St., SCEN 601 Fayetteville, AR, 72701.

## Abstract

All organisms experience stress and must rapidly respond to changing conditions. Thus, cells have evolved sophisticated rapid-response mechanisms such as post-translational protein modification to rapidly and reversibly modulate protein activity. One such post-translational modification is reversible lysine acetylation, where proteomic studies have identified thousands of acetylated proteins across diverse organisms. While the sheer size of the ‘acetylome’ is striking, the function of acetylation for the vast majority of proteins remains largely obscure. Here, we show that global acetylation plays a previously unappreciated role in the heat shock response of *Saccharomyces cerevisiae.* We find that dysregulated acetylation renders cells heat sensitive, and moreover, that the acetylome is globally remodeled during heat shock over time. Using quantitative acetyl-proteomics, we identified ∼400 high-confidence acetyl marks across ∼200 proteins that significantly change in acetylation when cells are shifted to elevated temperature. Proteins with significant changes in lysine acetylation during heat shock strongly overlap with genes induced or repressed by stress. Thus, we hypothesize that protein acetylation augments the heat shock response by activating induced proteins and inactivating repressed proteins. Intriguingly, we find nearly 40 proteins with at least two acetyl marks that significantly change in the opposite directions. These proteins are strongly enriched for chaperones and ribosomal proteins, suggesting that these two key processes are coordinately regulated by protein acetylation during heat shock. Moreover, we hypothesize that the same type of activating and inactivating marks that exist on histones may be a general feature of proteins regulated by acetylation. Overall, this work has identified a new layer of post-translational regulation that likely augments the classic heat shock response.

## Introduction

All organisms experience stress and must rapidly respond to changing environmental conditions. When the environment changes slowly, transcriptional responses are sufficient for altering protein levels to maintain the cellular milieu. On the other hand, rapid change necessitates an immediate response—the cell cannot afford to wait for newly synthesized stress defense proteins to meet acute demands. For these critically important proteins, cells maintain high enough basal or intrinsic levels of expression to weather short-term challenges. However, maintaining certain proteins in a constantly active state can be energetically wasteful (e.g., for ATP-consuming enzymes), or can lead to inappropriate activation of various cellular pathways. Thus, cells have evolved post-translational protein modifications (PTMs) to rapidly and reversibly modulate the activity of pre-existing proteins.

One such important PTM is reversible lysine acetylation, which was originally discovered as a histone modification over 50 years ago [1], and was immediately hypothesized to play a regulatory role. Histone acetylation is now a clearly established feature of eukaryotic gene regulation [2, 3], including under stressful conditions [4–8]. The observation that the human transcription factor and tumor suppressor p53 was regulated by reversible lysine acetylation [9] paved the way for the discovery that many other transcription factors are acetylation targets [10, 11].

Acetylation was thought to solely regulate transcriptional processes until the surprising discovery that acetylation regulates an enzyme—acetyl-CoA synthetase (Acs) [12]. Acs acetylation was found to be conserved from bacteria through humans [13–15], suggesting that the acetylation of enzymes and other non-transcriptional proteins may be widespread. Indeed, subsequent global proteomic studies have identified thousands of acetylated proteins in diverse organisms, rivaling phosphorylation in terms of the number of potential targets [16–23]. Several striking conclusions have emerged from these studies. First, many acetylation targets are shared in organisms ranging from bacteria to humans, suggesting that the regulatory functions of acetylation are likely well conserved. Second, the majority of acetylation targets are non-nuclear, including mitochondrial, ribosomal, and metabolic proteins, suggesting that acetylation may regulate many diverse processes in the cell.

The discovery of a large ‘acetylome’ raises questions about how global acetylation changes are regulated as well as their functional consequences. In terms of regulation, the key ‘writers’ and ‘erasers’ of lysine acetylation are the highly conserved lysine acetyltransferases (KATs) and lysine deacetylases (KDACs). While we have a limited understanding of how global protein acetylation patterns are regulated by KATs and KDACs, the metabolic state of the cell likely plays a major role. First, the substrate of KATs—acetyl-CoA—is a key central metabolite whose levels likely regulate KAT activity [24–26]. Second, one key class of KDACs—the sirtuins—use NAD^+^ as a substrate and generate nicotinamide as one of their products [27, 28]. The NAD^+^ to NADH ratio likely modulates sirtuin activity [29–31], and nicotinamide itself is a potent inhibitor [32, 33]. Finally, KATs and KDACs are generally members of large multi-subunit protein complexes, and subunit composition and modification can influence catalytic activities [34–37], protein substrate recognition [38, 39], and subcellular localization [40].

The large size of the acetylome also raises questions about the functions of acetylation on a global scale, and their regulatory role during environmental perturbation. We have obtained several lines of evidence suggesting that protein acetylation may play a key role in regulating the heat shock response in the model yeast *Saccharomyces cerevisiae.* First, yeast protein targets of acetylation from global proteomic surveys are strongly enriched for stress defense proteins, with the response to heat and protein folding being particularly noteworthy [16, 41]. Second, there is growing evidence that acetylation can regulate the activity of key stress defense proteins in yeast and other organisms. For example, the mammalian and human heat shock factor HSF1 that coordinates the transcriptional response to heat shock is regulated by reversible lysine acetylation [42, 43]. Moreover, acetylation has been shown to regulate the direct ‘first responder’ heat shock chaperone proteins, impacting activity and/or client interactions [44–47]. Indeed, reversible lysine acetylation has been hypothesized to be a bedrock component of the ‘chaperone code’ [48, 49], which similar to its namesake histone code, is a complex array of PTMs that may regulate chaperone activity, client specificity, and localization.

Despite these connections of acetylation to stress defense, there have been few studies examining whether the acetylome is systematically remodeled during stress responses in any organism. Acetylation changes during salt stress have been observed in the bacterium *Bacillus altitudinis* [50], with differential acetylation of proteins involved in osmolyte biosynthesis and antioxidant defense being noteworthy. The effects of heat shock on the acetylome have been explored in two marine invertebrates: Pacific oyster *Crassostrea gigas* [51] and sea cucumber *Apostichopus japonicus* [52]. Both marine studies found differential acetylation during heat shock of chaperone proteins and metabolic enzymes, though the lack of normalization to changes in total protein abundance warrants cautious interpretation. As such, a comprehensive analysis of how the acetylome is dynamically remodeled during stress in a genetically tractable eukaryotic model system has been lacking.

Here, we show that acetylome remodeling adds an important layer to the heat shock response of *Saccharomyces cerevisiae.* We find that inhibition of deacetylation impacts cellular fitness during heat stress, which correlates with extensive global acetylome remodeling during heat shock. Using quantitative acetyl-proteomics, we identified ∼400 high-confidence acetyl marks across ∼200 proteins that significantly change in acetylation when cells are shifted to elevated temperature. Proteins with significant changes in lysine acetylation during heat shock strongly overlap with genes induced or repressed by heat stress. Thus, we hypothesize that protein acetylation augments the heat shock response by activating induced proteins and inactivating repressed proteins. Nearly 40 proteins display both increasing and decreasing acetylation marks on different residues, suggesting precise regulatory control by acetylation that is evocative of the histone and chaperone codes. Through analysis of metabolites, enzyme complexes, and protein localization, we identify multiple mechanisms that could contribute to acetylome remodeling during heat shock. Overall, our results reveal protein acetylation as a previously unappreciated layer of global post-translational control that likely augments the classic heat shock response.

## Results and Discussion

### Acetylation is required for full acquisition of thermotolerance in the absence of protein synthesis

Given the substantial circumstantial evidence linking the acetylome to stress defense [16, 41], and heat shock in particular, we sought to investigate how acetylation might contribute to heat stress defense independently of its well-established transcriptional role [5–8, 42, 43, 53–55]. One approach to isolate transcription-independent processes is to include the protein synthesis inhibitor cycloheximide concurrently to assessing stress defense phenotypes. However, because growth requires nascent protein synthesis, this precludes the use of cycloheximide when assessing growth phenotypes, which are most frequently used to assess cellular fitness during stress. Fortunately, previous studies examining a phenomenon called acquired thermotolerance [56–58], where cells exposed to mild heat stress become resistant to an otherwise lethal severe dose of heat, found that this phenomenon can occur in the absence of protein synthesis [59–61]. Thus, we sought to examine the effects of perturbing global acetylation on the cell’s ability to acquire thermotolerance in the absence of protein synthesis.

Consistent with previous studies [59–61], we observed a large increase in survival at 45°C when cells were first pretreated with a mild 37°C heat shock compared with control cells that were maintained at 25°C (Figure 1B and 1C). Treatment with cycloheximide caused only a minor, statistically insignificant decrease in thermotolerance acquisition, confirming that acquired thermotolerance can occur largely independently of nascent protein synthesis at this temperature dose. Intriguingly, we found that treatment with a cocktail of deacetylase inhibitors (trichostatin A [62] and nicotinamide [63]) led to modest, but significantly impaired acquired thermotolerance in the absence of protein synthesis (Figure 1). While nicotinamide does have potential sirtuin-independent effects [64], the impairment of acquired thermotolerance in the presence of KDAC inhibitors is also consistent with acetylation playing a role in regulating heat defense. We note that acquired thermotolerance is not completely lost in the presence of KDAC inhibitors, suggesting that other post-translational modifications may also play a role. However, another (non-mutually exclusive) possibility is that both acetylation and deacetylation are important for acquired thermotolerance (e.g., acetylation activates a specific set of stress defense proteins, while deacetylation activates a second set), but on net, deacetylation plays a larger role. Because the effect was modest, we examined whether treatment of cells with KDAC inhibitors alone would affect other heat stress phenotypes. We did not observe a defect in acquired thermotolerance when cells were treated with KDAC inhibitors alone, suggesting that the modest effect of KDACs on acquired thermotolerance only manifests in the absence of nascent protein synthesis (Figure 1B). In contrast, we did observe a very strong effect of KDAC inhibition on growth at 40°C (Figure 1D), suggesting that properly regulated protein (de)acetylation is important for fitness during heat stress. Because of this phenotypic link, we next sought to explore globally how the acetylome may be remodeled during heat shock.

**Figure 1.**
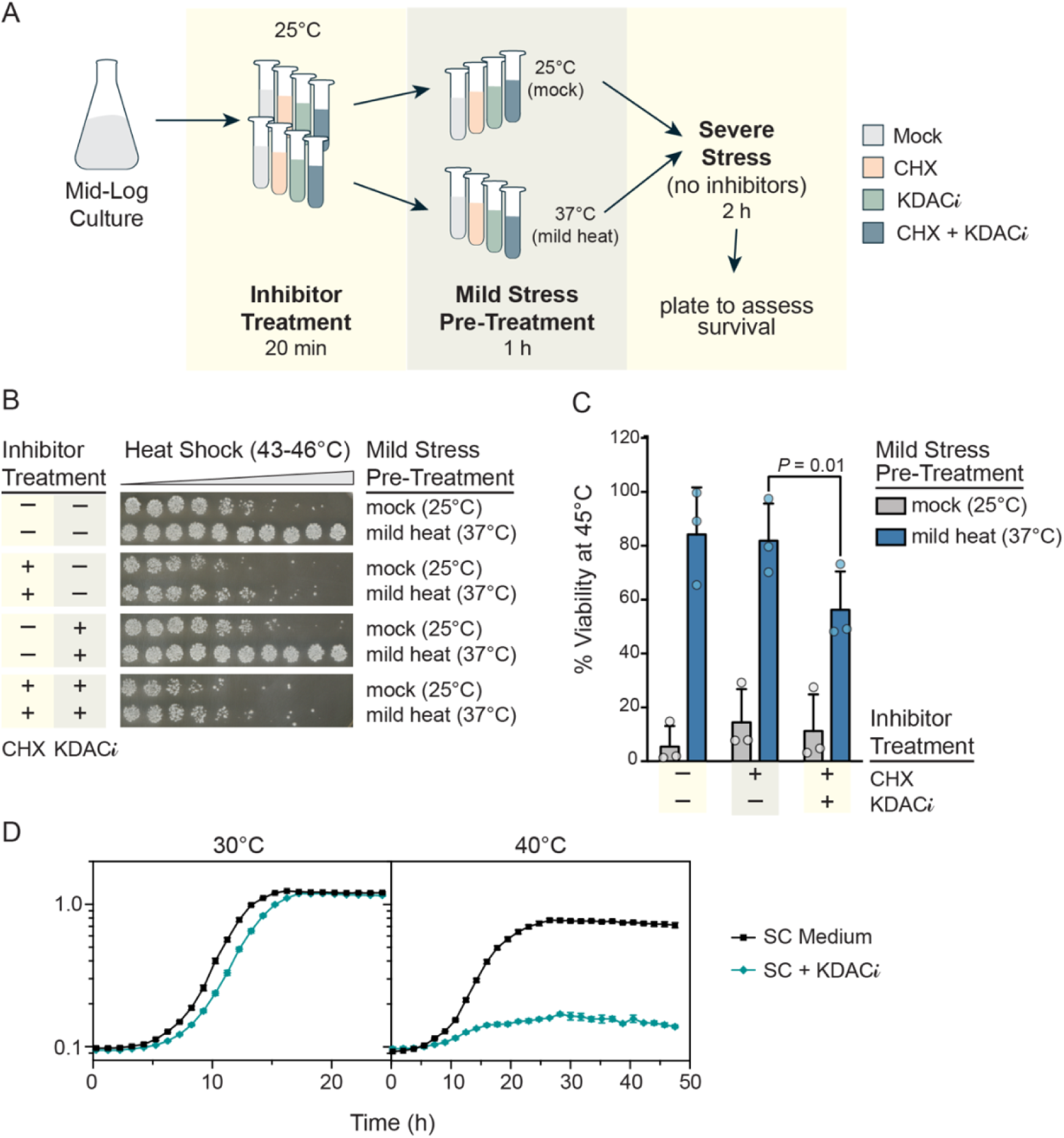
Acquired thermotolerance in the absence of protein synthesis and growth at 40°C requires lysine deacetylase activity. (A) Schematic of acquired thermotolerance assay. Cells received either pre-warmed 25°C media (mock) or 37°C media (mild heat) for 1 h, before being shifted to severe heat for 1 hour. Cycloheximide (CHX; 10 μg / ml) was used as a protein synthesis inhibitor, and trichostatin A (10 μM) and nicotinamide (5 mM) were used as lysine deacetylase inhibitors (KDAC*i*) Inhibitor-treated cells were exposed 20 min before and throughout mock or mild heat treatment. (B) Representative acquired thermotolerance assays performed using a thermal cycler with a gradient. (C) Acquired thermotolerance assay conducted with a single severe heat dose (45°C) and quantified via CFU counting. Error bars denote the standard deviation of biological triplicates, with the *P*-value being calculated using a paired *t*-test. (D) Growth assays of cells grown in minimal media in the presence or absence of KDAC*i* at either 30°C or 40°C. Error bars denote the standard error of 3 biological replicates.

### Heat shock triggers extensive remodeling of the yeast acetylome

To understand the dynamics of potential acetylome remodeling during heat shock, we performed quantitative acetyl-proteomics across a four-hour 25°C to 37°C heat shock time course in biological triplicate (see Methods). High-confidence acetyl-lysine peptides were defined as being quantified in all TMT channels (i.e., all time points) in all 3 biological replicates. In total, this resulted in quantification of 1,473 acetyl-lysine residues across 598 total proteins (Table S3). Consistent with previous yeast acetyl-proteomic surveys [41], the acetylated proteins we identified were enriched for similar functional categories (Table S4).

To identify proteins whose acetylation state significantly changed during heat shock, we first calculated log_2_ fold changes in acetyl-peptide abundance at each time point relative to the unstressed control. Then, to control for any underlying changes in total protein abundance during heat shock, we further normalized to log_2_ fold changes in protein abundance (i.e., acetylation changes for any given protein were normalized to changes in underlying total abundance for that same protein). Finally, we used statistical significance testing on the resulting normalized acetyl-peptide log_2_ fold changes with a false discovery rate (FDR) threshold of <0.05 to identify acetyl-peptides with significant abundance changes. In total, we identified 387 residues across 207 proteins that showed a significant change in acetylation in at least one time point compared to the unstressed control (Figure 2A and Table S3).

**Figure 2.**
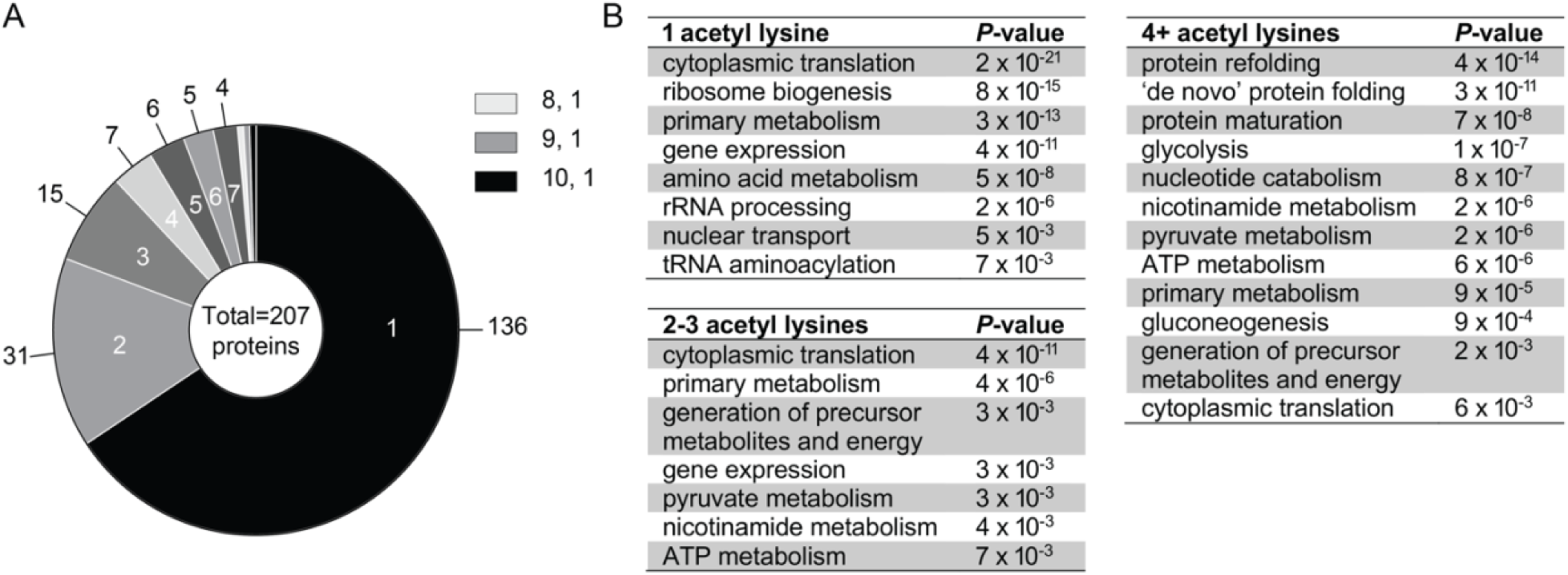
Heat shock remodels the yeast acetylome, with a subset of proteins experiencing extensive remodeling. (A) The pie chart depicts the number of acetyl lysine peptides with significant differential abundance for at least one time point post 25°C to 37°C heat shock (FDR < 0.05). (B) Functional enrichments of proteins with the indicated number of acetyl lysine residues with significant differential abundance.

We next examined the properties of proteins with acetyl lysines that changed in abundance. Proteins with at least one significantly changing acetyl lysine residue were notably enriched for those involved in cytoplasmic translation (*P =* 1 x 10^-38^), ribosome biogenesis (*P =* 2 x 10^-19^), gene expression (*P =* 7 x 10^-19^), and protein refolding (*P =* 3 x 10^-13^), suggesting that these processes may be regulated by reversible lysine acetylation during heat shock. Intriguingly, while the majority (66%) of proteins with differential acetyl lysine abundance during heat shock had a single changing residue, a substantial fraction had multiple changing sites. The translational elongation factor eEF1A (Tef1) showed the most extensive dynamic acetylation remodeling, with 10 distinct sites changing in abundance during heat shock. This extensive modification is intriguing in light of eEF1A’s known role in heat defense. In mammals, eEF1A plays a key role in the heat shock response by recruiting the main heat shock transcription factor HSF1 to heat-shock protein promoters and facilitating nuclear export of the nascent mRNAs [65]. In both mammals and yeast, lack of eEF1A renders cells sensitive to heat stress [65, 66]. More broadly, the functions of proteins with multiple dynamic acetyl sites were distinct compared to the proteins with a single changing site (Figure 2B). While proteins with a single dynamic site were strongly enriched for cytoplasmic translation and gene expression, proteins with 4 or more dynamic acetyl lysine residues showed much stronger enrichment for protein folding functions (Figure 2B). This pattern suggests that dynamic regulation of multiple acetylation sites may be particularly important for fine-tuning chaperone function during heat stress.

Analysis of acetylation dynamics revealed that peak remodeling occurred at 90 minutes, largely coinciding with the timing of the peak translational response (Figure 3). This was a surprising result, as we originally hypothesized that acetylome remodeling would precede changes in protein abundance, similar to what has been observed for phosphorylation dynamics during heat shock [67]. However, several lines of evidence suggest that the observed patterns of acetylation are not simply following protein level changes. First, despite normalization to underlying changes in protein abundance, the magnitude of acetylation abundance changes were generally larger than protein-level changes. Second, we observed that 69 out of 208 proteins (33%) showed significant changes in protein acetylation levels despite no change in underlying protein levels over the course of the experiment, including the abovementioned Tef1 with the most dynamic acetylation sites. Finally, we identified dozens of proteins that displayed both increasing and decreasing acetyl lysine abundance on different residues, strongly suggesting precise effects (Figure 4). Altogether, our data suggest that acetylome remodeling during heat shock does not simply reflect protein-level changes, but instead represents a distinct regulatory program.

**Figure 3.**
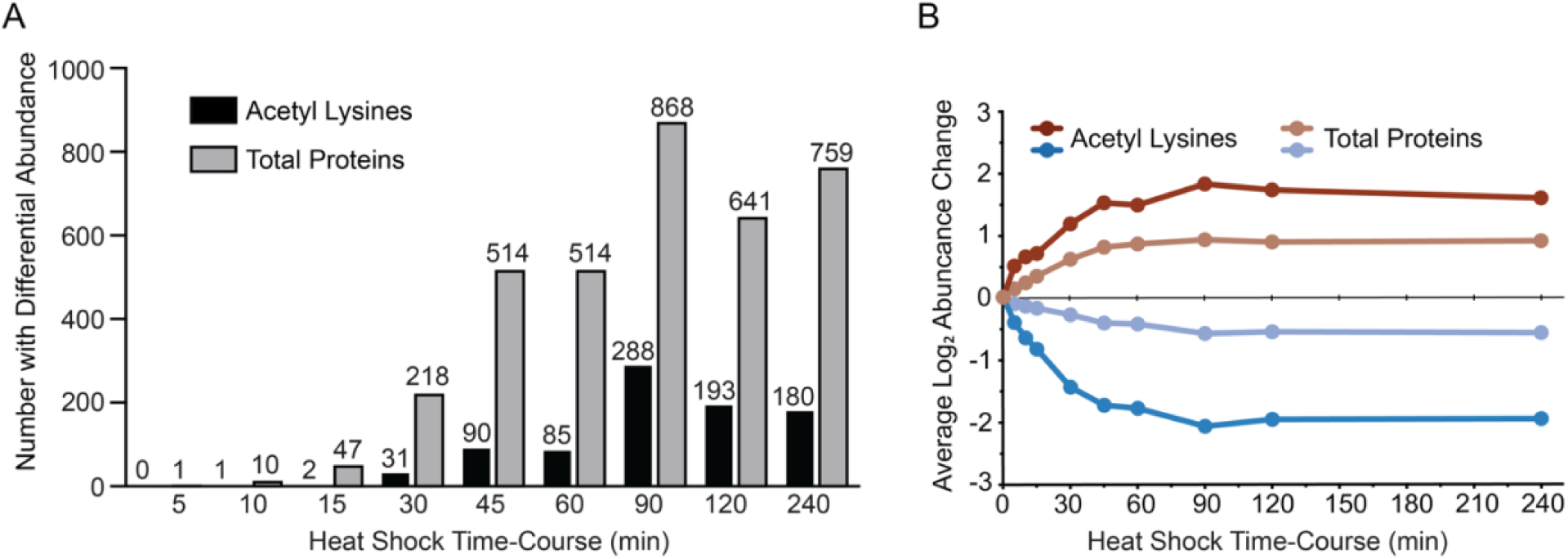
Peak acetylome remodeling occurs at 90 minutes post-heat shock. (A) The bars depict the number of acetyl-lysine peptides or total proteins with significantly differential abundance (FDR < 0.05) at each time point post-heat shock (25°C to 37°C) relative to an unstressed control. (B) The average log_2_ fold change at each time point for all acetyl lysine peptides or total proteins experiencing significant differential abundance in at least 3 time points (FDR < 0.05).

**Figure 4.**
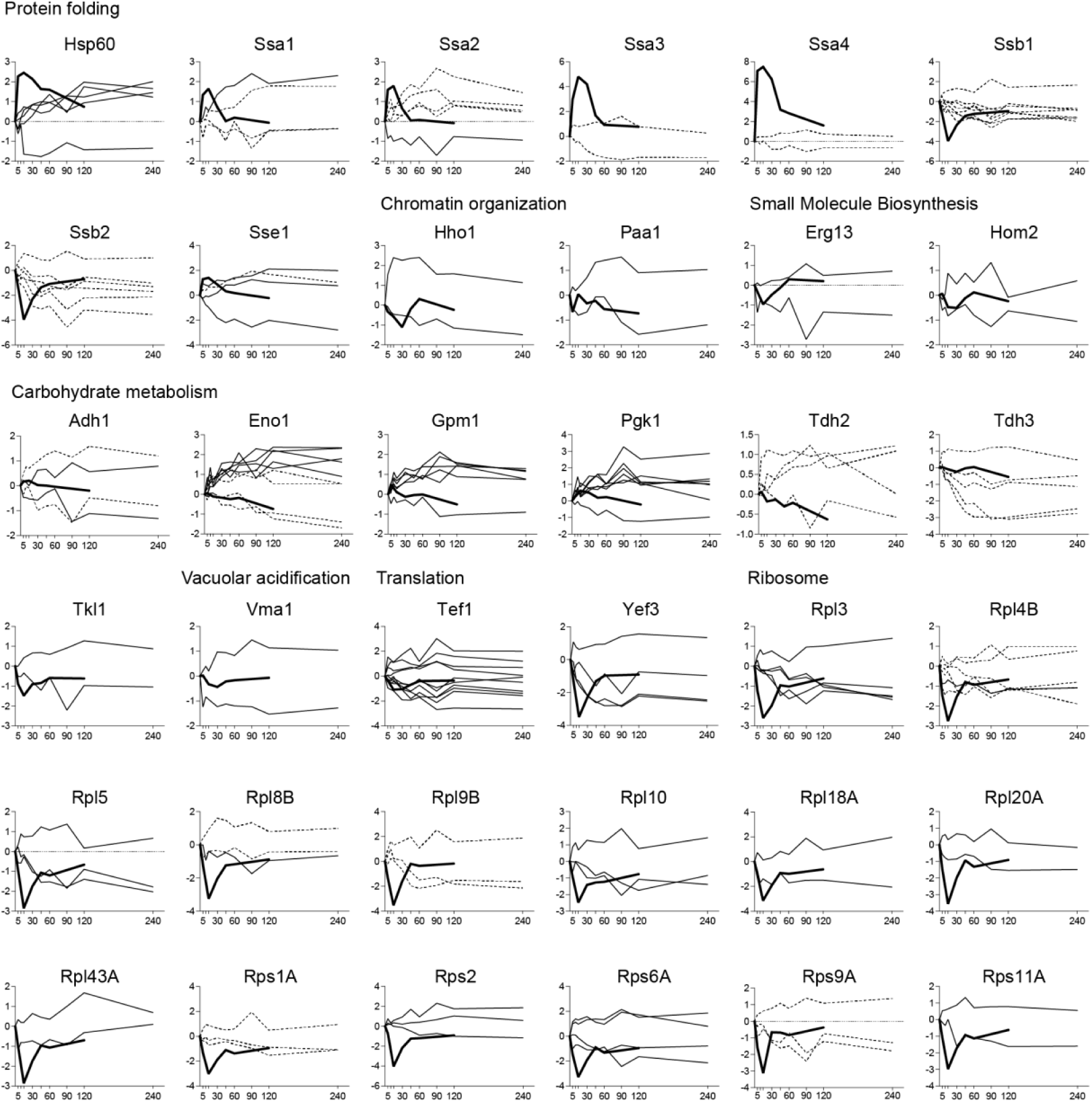
Proteins that experience both increasing and decreasing acetyl residues. For each protein, the thin lines represent the average log_2_ fold change (y-axis) of one acetyl-lysine residue over the 240-min 25°C to 37°C time course (x-axis). The thick lines represent mRNA log_2_ fold changes during a 25°C to 37°C heat shock time course from [123]. Proteins are grouped based on their annotated functions. Note that some acetyl-peptides (denoted with dotted lines) can be ambiguously assigned to two or more proteins (generally paralogs, see Table S3 for the full list). Twenty-two proteins have lysines that unambiguously increased or decreased in acetylation during heat shock.

### Acetylation dynamics during heat shock reveal distinct functional patterns

To better understand temporal patterns of acetylome remodeling during heat shock, we performed hierarchical clustering of the 387 significantly changing acetyl sites. Proteins showing increased acetylation during heat shock were strongly enriched for functions related to protein folding and refolding, carbohydrate metabolism, and ATP synthesis (Figure 5A and Table S4). Conversely, proteins with decreasing acetylation were enriched for translation, ribosome biogenesis, and nitrogen metabolism (Figure 5A and Table S4). The functions of these proteins with changing acetyl lysine residues strikingly mirrors the canonical transcriptional response to heat shock [68], where heat shock protein chaperones and stress defense proteins are induced, and translational-related genes are repressed. To directly test whether the top genes regulated transcriptionally during the heat shock response are also subject to dynamic acetylation, we examined whether the genes most strongly transcriptionally induced or repressed during heat shock were more likely to encode proteins subject to dynamic acetylation. To do this, we pulled the top 25 transcriptionally induced or repressed genes during heat shock from [69] that also had at least one observed acetyl-lysine in our dataset. Of those top 25 induced genes, 16/25 of the encoded proteins had a dynamic acetylation event (Table 1), which was more than expected by chance considering the total of proteins with dynamic acetylation in our dataset (207) versus total acetylated proteins (598) (*P* = 0.002, 1.8-fold over-enrichment, Fisher’s exact test). Likewise, 15/25 top repressed genes encoded proteins with dynamic acetylation (Table 2), which was also a significant enrichment (*P* = 0.007, 1.7-fold over-enrichment, Fisher’s exact test). Thus, we hypothesize that protein acetylation serves as an additional layer of regulation that augments these transcriptional and translational changes—activating proteins involved in stress defense while suppressing proteins involved in growth-related processes.

**Figure 5.**
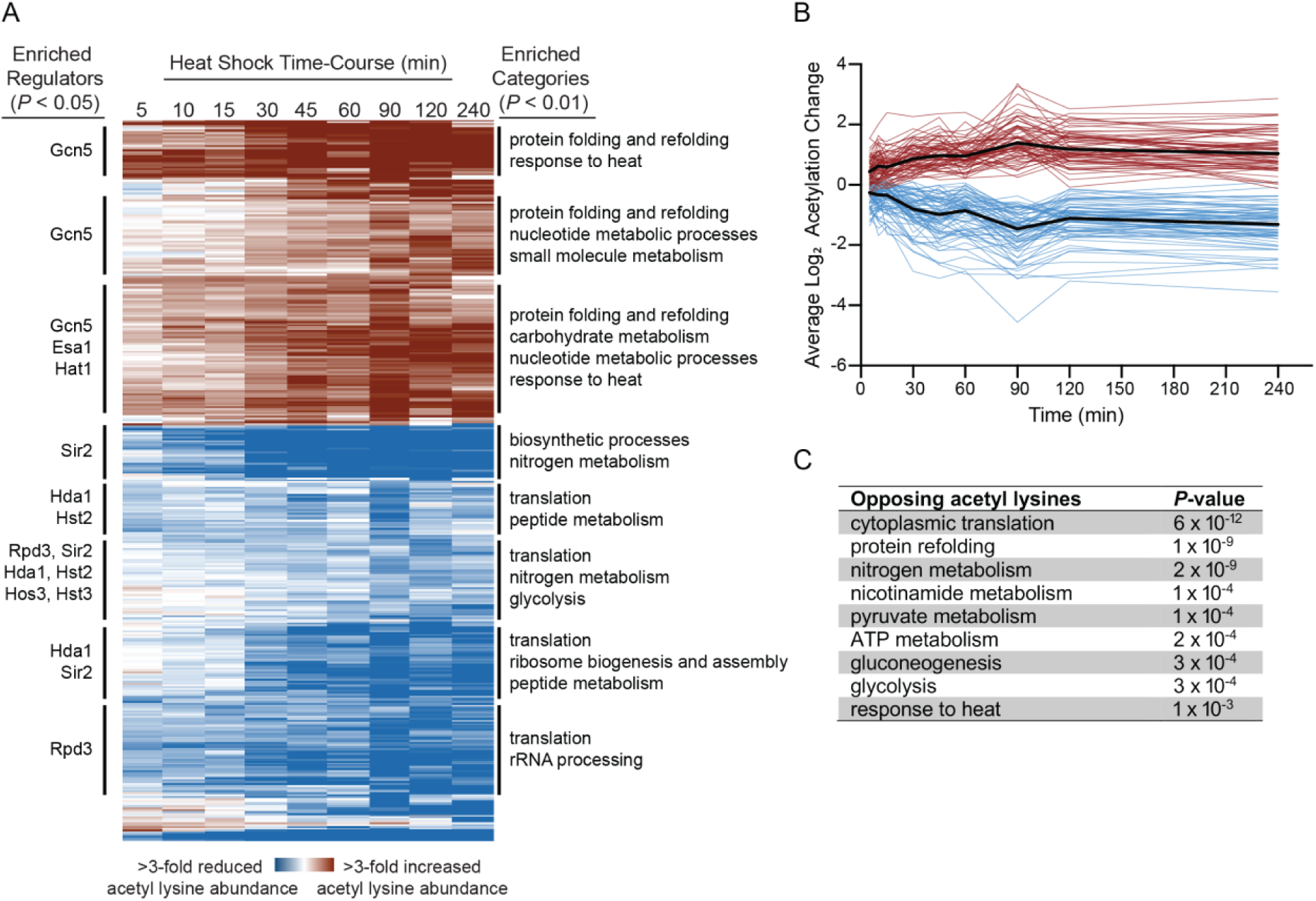
Heat shock induces distinct temporal patterns of acetylome remodeling that correlate with specific cellular functions. (A) The heat map depicts hierarchical clustering of log_2_ fold changes for acetyl lysine peptides with significant changes in abundance during heat shock (25°C to 37°C) for at least one time point (FDR < 0.05). Each row depicts a single acetyl lysine peptide, and each column depicts the average abundance change during heat shock relative to the unstressed control at the indicated time points. Red denotes increased lysine acetylation and blue denotes decreased lysine acetylation as indicated in the key. Significant regulatory enrichments (KATs and KDACs) are annotated to the left, and functional enrichments are annotated to the right (Bonferroni-corrected P < 0.01). (B) Average log_2_ fold changes in acetyl lysine abundance following heat shock for 36 proteins showing opposing acetylation state changes on different residues. (C) Functional enrichments for the 36 proteins with opposing acetyl marks, with the caveat that some acetyl marks cannot be unambiguously assigned (Figure 4 and Table S3).

**Table 1.** Heat shock-induced genes are enriched for proteins with dynamic acetylation.

| Gene ID | Gene Name | log<sub>2</sub>FC (mRNA)<sup>1</sup> | Total AcK Sites | Increasing<sup>2</sup> | Decreasing<sup>2</sup> | Unchanged<sup>2</sup> |
| --- | --- | --- | --- | --- | --- | --- |
| YER103W | SSA4* | 7.20 | 7 | 1 | 1 | 5 |
| YBL075C | SSA3* | 5.09 | 6 | 1 | 1 | 4 |
| YLL026W | HSP104 | 4.62 | 9 | 6 | 0 | 3 |
| YPL240C | HSP82* | 4.32 | 21 | 7 | 0 | 14 |
| YGR142W | BTN2 | 3.76 | 1 | 0 | 0 | 1 |
| YBR169C | SSE2* | 3.61 | 1 | 1 | 0 | 0 |
| YCL040W | GLK1 | 3.39 | 1 | 0 | 0 | 1 |
| YLR216C | CPR6 | 3.34 | 4 | 1 | 0 | 3 |
| YJL052W | TDH1* | 3.25 | 7 | 2 | 0 | 5 |
| YAL005C | SSA1* | 3.16 | 13 | 2 | 2 | 9 |
| YCL042W | YCL042W | 3.04 | 1 | 1 | 0 | 0 |
| YLR438W | CAR2 | 3.00 | 1 | 0 | 0 | 1 |
| YOR027W | STI1 | 2.68 | 9 | 5 | 0 | 4 |
| YMR303C | ADH2* | 2.65 | 3 | 0 | 1 | 2 |
| YKL035W | UGP1 | 2.55 | 3 | 2 | 0 | 1 |
| YOR185C | GSP2* | 2.47 | 5 | 0 | 1 | 4 |
| YGR254W | ENO1* | 2.44 | 16 | 7 | 2 | 7 |
| YNL007C | SIS1 | 2.27 | 1 | 0 | 0 | 1 |
| YMR120C | ADE17 | 2.18 | 1 | 0 | 0 | 1 |
| YGL062W | PYC1 | 2.14 | 1 | 0 | 0 | 1 |
| YLR259C | HSP60 | 2.10 | 7 | 4 | 1 | 2 |
| YJR148W | BAT2 | 2.07 | 1 | 0 | 0 | 1 |
| YCL064C | CHA1 | 1.98 | 2 | 0 | 0 | 2 |
| YJL034W | KAR2 | 1.97 | 4 | 1 | 0 | 3 |
| YPR035W | GLN1 | 1.85 | 2 | 0 | 0 | 2 |

<sup>1</sup> The top 25 most strongly heat shock-induced genes from [69] that encode a protein with an observed acetyl-lysine are listed. Log<sub>2</sub> fold changes represent relative mRNA abundance 20-minutes post 25°C to 37°C heat shock relative to an unstressed control (cell grown at 25°C).
<sup>2</sup> For corresponding protein, acetyl-lysine sites were classified as increasing (positive log<sub>2</sub> fold change) or decreasing (negative log<sub>2</sub> fold change), based on the majority direction of change at significant time points (FDR < 0.05) post 25°C to 37°C heat shock. For
\* denotes that one or more significantly changing acetyl-marks are ambiguous (see Table S3 for a full list of ambiguously mapped acetyl marks)

**Table 2.**
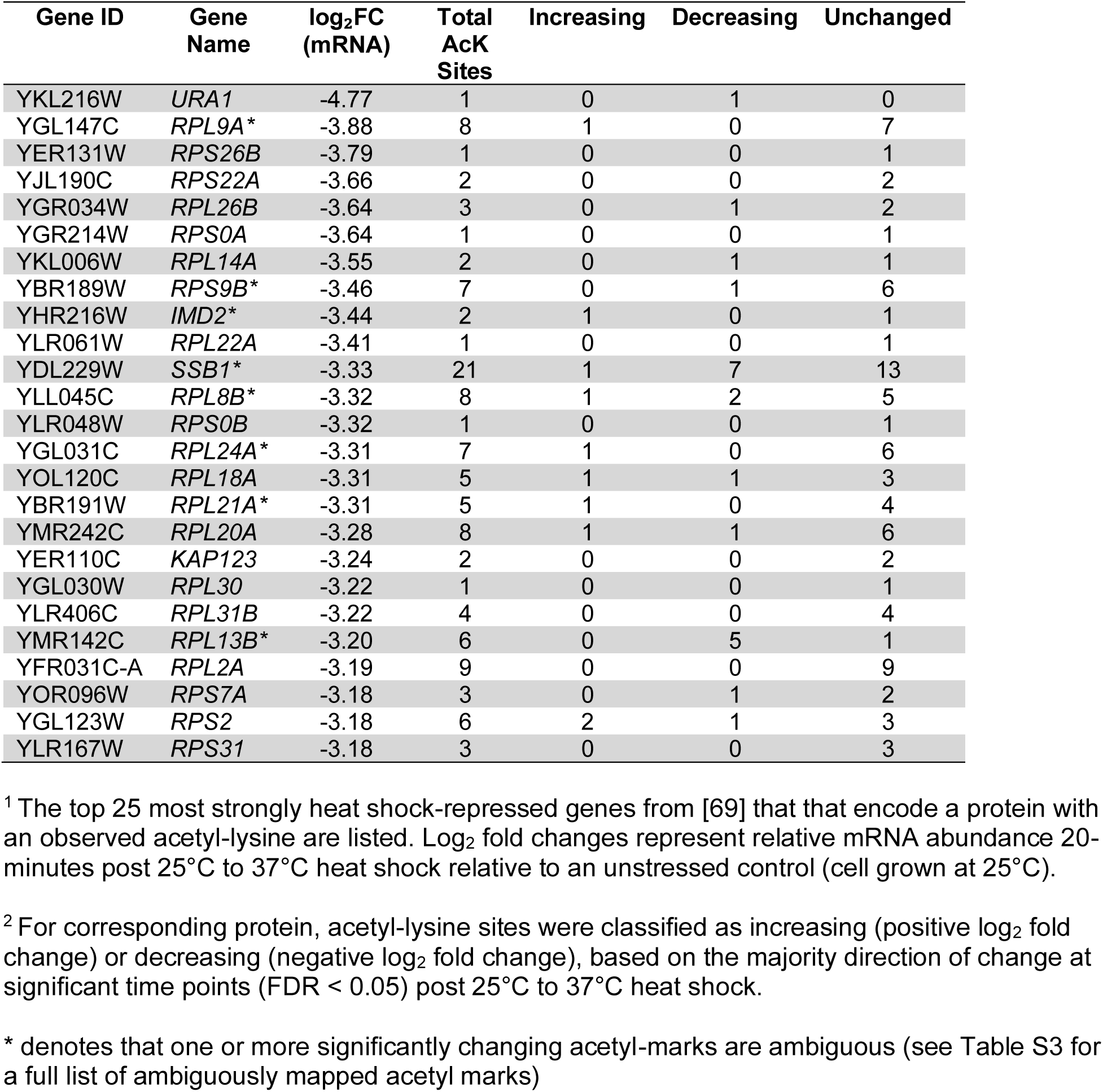
Heat shock-repressed genes are enriched for proteins with dynamic acetylation.

Intriguingly, we identified 36 proteins that displayed both increasing and decreasing acetylation on different lysine residues (Figure 4 and Figure 5B). These proteins were strongly enriched for chaperones and ribosomal proteins (Figure 5C), suggesting that acetylation may play a particularly important aspect of regulation of these two key processes during heat shock. We note that some acetyl-peptides map to more than one paralog and cannot be unambiguously assigned to a single protein (see Table S3 for the full list), with 22 of the 36 proteins having at least one unambiguous opposing pair of marks. We note potential parallels to the histone code, where the same type of activating and inactivating marks that exist on histones may be a general feature of proteins regulated by acetylation. Indeed, different constellations of PTMs have been hypothesized to form a fundamental part of the ‘chaperone code’ [48, 49] to fine tune their function, substrate specificity, and interaction networks. Our discovery of differential acetylation on multiple chaperones suggests that acetylome remodeling during heat shock may be an important aspect of this code, fine tuning chaperone activity during heat stress. Additionally, there is evidence that heat stress triggers adaptive phase separation [70]. Since protein acetylation can regulate phase separation behavior [71], the complex patterns of chaperone acetylation we observe could help regulate formation and/or activities of stress-induced biomolecular condensates. Similarly, the complex acetylation patterns on ribosomal proteins might help orchestrate the dramatic translational reprogramming that occurs during stress [72–74], potentially fine-tuning which mRNAs continue to be translated. Together, these results suggest that acetylation provides an additional layer of post-translational control that complements and enhances the classical heat shock response.

### Genetic evidence links Sti1 acetylation to thermotolerance

To directly test whether acetylation changes affect protein function during stress, we focused on Sti1, an Hsp70 and Hsp90 co-chaperone. Sti1 was chosen because unlike other potential chaperones or co-chaperones with dynamic acetyl sites, acetylation has not been shown to date to affect Sti1 function. During our acetyl-proteomic time course, Sti1 showed increased acetylation at residues K54 and K337 during heat shock. We created mimetic mutants either preventing acetylation (arginine substitutions K54R and/or K337R) or mimicking constitutive acetylation (glutamine substitutions K54Q and/or K337Q), as well as non-conservative alanine substitutions to determine whether the lysines are functionally important (K54A and/or K337A). We introduced the mutant alleles on a plasmid in a *sti1*Δ deletion strain that was unable to grow at 40°C (Figure 6). For both K54 and K337, the unacetylatable arginine mutants grew similarly to wild-type strain, while the acetyl-lysine mimetic mutants showed growth defects similar to the alanine substitutions (Figure 6), suggesting that lysine acetylation at those sites reduces Sti1 function. Combining both acetyl-lysine mimetic mutations (K54Q and K337Q) did not result in an additive fitness defect beyond the individual single mutations.

**Figure 6.**
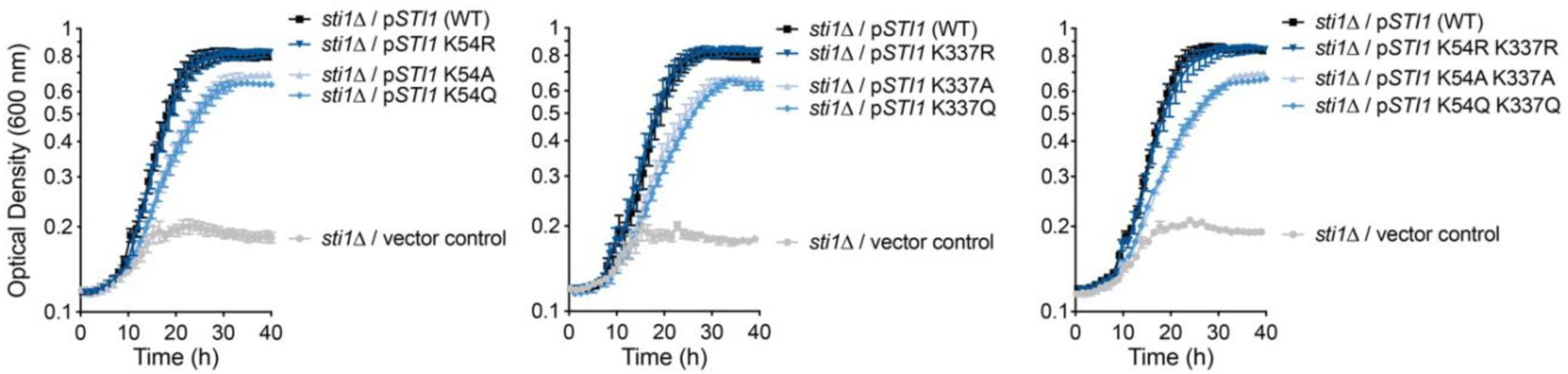
Genetic evidence of Sti1 regulation by reversible lysine acetylation during heat shock. Growth assays were performed at 40°C comparing wild-type *STI1* with mutants mimicking either constitutive acetylation (K54Q and/or K337Q) or unacetylatable lysine (K54R and/or K337R). Alanine substitutions were included as controls to determine the functional importance of each lysine. Error bars denote the standard error of 6 biological replicates.

The impaired growth of cells expressing constitutively ‘acetylated’ Sti1 could reflect the importance of dynamic regulation of acetylation on Sti1 function (vs the static state of the mimetic mutants). Sti1 serves as a scaffold protein that helps coordinate the activities of Hsp70 and Hsp90 through distinct tetratricopeptide repeat (TPR) domains [75–77]. K54 lies within the TPR1 domain that mediates Hsp70 binding, while K337 is in the TPR2A domain responsible for Hsp90 interactions [77]. The location of these acetylation sites in domains mediating key protein-protein interactions suggests acetylation could modulate how Sti1 coordinates these essential chaperone systems during heat shock, and is consistent with the emerging concept of the chaperone code [48, 49]. It is also possible that heat stress intrinsically increases Sti1 acetylation to reduce function, and the action of KDACs are a necessary counterbalance to preserve function. Future biochemical studies examining how acetylation affects Sti1’s function and/or interactions with Hsp70 and Hsp90 will help distinguish these possibilities.

### Possible regulators of acetylome remodeling during heat shock

The extensive acetylome remodeling during heat stress raised important questions about their regulation. Several non-mutually exclusive possibilities could regulate these dynamics. At a global level, changes in key metabolites that are substrates or inhibitors of KATs and/or KDACs could be responsible for acetylation dynamics. More specific regulation could occur through changes in individual KAT/KDAC abundance, remodeling of membership of KAT/KDAC complexes, and/or changes in KAT/KDAC subcellular localization. To explore these possibilities, we systematically investigated each of these potential regulatory mechanisms.

We first examined changes in key metabolites that can influence protein acetylation: acetyl-CoA, free CoA, nicotinamide, and the NAD^+^ to NADH ratio. Acetyl-CoA serves as the acetyl donor for all known KAT reactions [24–26], while CoA can act as a competitive inhibitor [78, 79]. During heat shock, cellular acetyl-CoA levels were reduced to approximately 50% of unstressed levels by 60 minutes, while CoA levels correspondingly increased by 50% at 30 minutes before recovering at 60 minutes (Figure 7). Nicotinamide (an inhibitor of sirtuins [32, 33].) levels mildly spiked at 5 minutes before decreasing by ∼40% at 30 minutes (Figure 7). The NAD^+^ to NADH ratio is thought to regulate sirtuin activity [29–31], and this ratio briefly spiked at 5 minutes before moderately decreasing at 60 minutes (Figure S1). Notably, all of these metabolite changes occurred earlier than the peak of acetylome remodeling at 90 minutes. This temporal disconnect suggests that global metabolite levels are not the primary drivers of the observed acetylation. While local concentrations of these metabolites in specific cellular compartments might still play a role [80], our data suggest that acetylome remodeling during heat shock is likely regulated by the action of specific KATs and/or KDACs.

**Figure 7.**
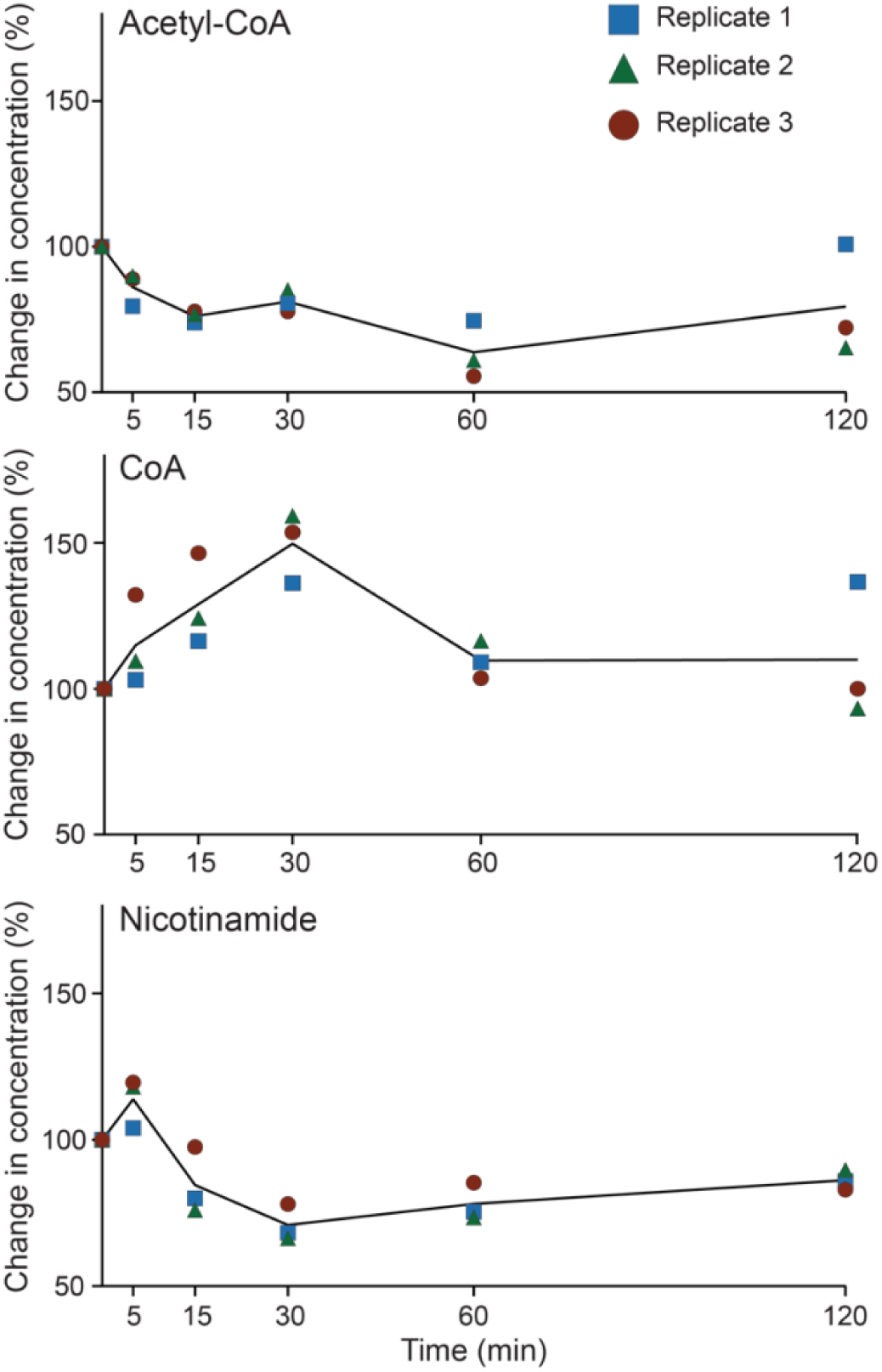
Key regulatory metabolites show rapid but largely transient changes during heat shock. Concentrations of (A) acetyl-CoA, (B) CoA, and (C) nicotinamide were determined across 120 minutes of 25°C to 37°C heat shock using LC-MS/MS.

To identify enzymes that might regulate these acetylation changes, we analyzed each cluster for enrichment of known physical interactions with KATs and KDACs. Each of the 8 clusters was enriched for interactions with at least one KAT or KDAC (Figure 4A), suggesting potential mechanistic links between specific enzymes and the observed patterns of acetylome remodeling. Clusters showing increased acetylation were strongly enriched for interactions with Gcn5, suggesting this KAT may play a central role in stress-induced protein acetylation. In contrast, clusters showing decreased acetylation were enriched for interactions with multiple KDACs including Sir2, Rpd3, She1, Hst2, Hst3, and Hos3. We performed motif analysis on acetyl-peptide sequences within each cluster to identify potential sequence specificity (Figures S2 and S3). While some clusters showed a weak enrichment for previously identified motifs, such as small amino acids at the -1 position (relative to the acetyl lysine) and KK pairs [81, 82], we did not identify strong cluster-specific sequence patterns. This could reflect the complexity of regulation of acetylome remodeling during heat stress, as we did observe enrichments for multiple KATs and/or KDACs for half of the clusters.

We also examined the total proteome data to identify changes in KAT and KDAC protein abundance over the heat shock time course. Across all 3 biological replicates, we were able to quantify relative abundance for 1 KAT (Hat1) and 4 KDACs (Hos3, Rpd3, She1, and Sir2) that were implicated in acetylation regulation via our clustering analysis. Among these enzymes, only Hat1 and Sir2 showed significant changes in abundance, both decreasing at 90 minutes post-heat shock. Notably, these changes in enzyme abundance did occur concurrently with the peak of acetylome remodeling, suggesting that altered Hat1 and Sir2 levels could play a part in a subset of acetylome remodeling. Because several KATs and KDACs were not represented in our total proteome data, we also examined our previous study examining the peak of the yeast transcriptional heat shock response [69]. Here, we additionally found one up-regulated KAT (*HPA2*), three down-regulated KATs (*GCN5, ELP3,* and *SAS3*), and two downregulated KDACs (*SAS2* and *RTT109*) (all at FDR < 0.01). In total, 9/23 (39%) of KATs and KDACs showed evidence of regulation by heat shock, suggesting that a potentially substantial fraction of global changes in acetylation during heat shock could be explained by changes in relative KAT and KDAC abundances. However, we also sought to determine KAT/KDAC relocalization, complex remodeling, and/or changes in substrate preference could also play a role in acetylome remodeling during heat shock.

Given that most KATs and KDACs function within multi-protein complexes that help determine their activities [34–37], substrate specificity [38, 39], and/or localization [40], we next investigated whether heat shock could induce changes in the subunit composition of these complexes. Using TAP-tagged KAT and KDAC strains, we performed co-immunoprecipitation experiments across the heat shock time course for potential regulatory KATs and KDACs (Figure 8 and Figure S4). Two KATs showed evidence of protein-protein interaction changes during heat shock. The essential KAT Esa1 is the catalytic subunit of the NuA4 complex [83], and Esa1 showed increased association with both the ribosomal protein Rpl12A and the vesicle membrane receptor Snc2 (Figure 8). While these proteins are not components of NuA4, their increased association with Esa1 suggests they could be specific substrates during heat shock (though notably these proteins were not identified as differentially acetylated in our acetyl-proteomics time course). The TFIID component Taf1 (thought to be a minor KAT [84]) showed an increased association with Tdh3 during heat stress (Figure 7). Tdh3 is a glyceraldehyde-3-phosphate dehydrogenase that is known to ‘moonlight’ and interact with Sir2 [85] and SAGA [86] to regulate transcription. Thus, it is possible that Taf1 plays a more significant role during heat shock.

**Figure 8.**
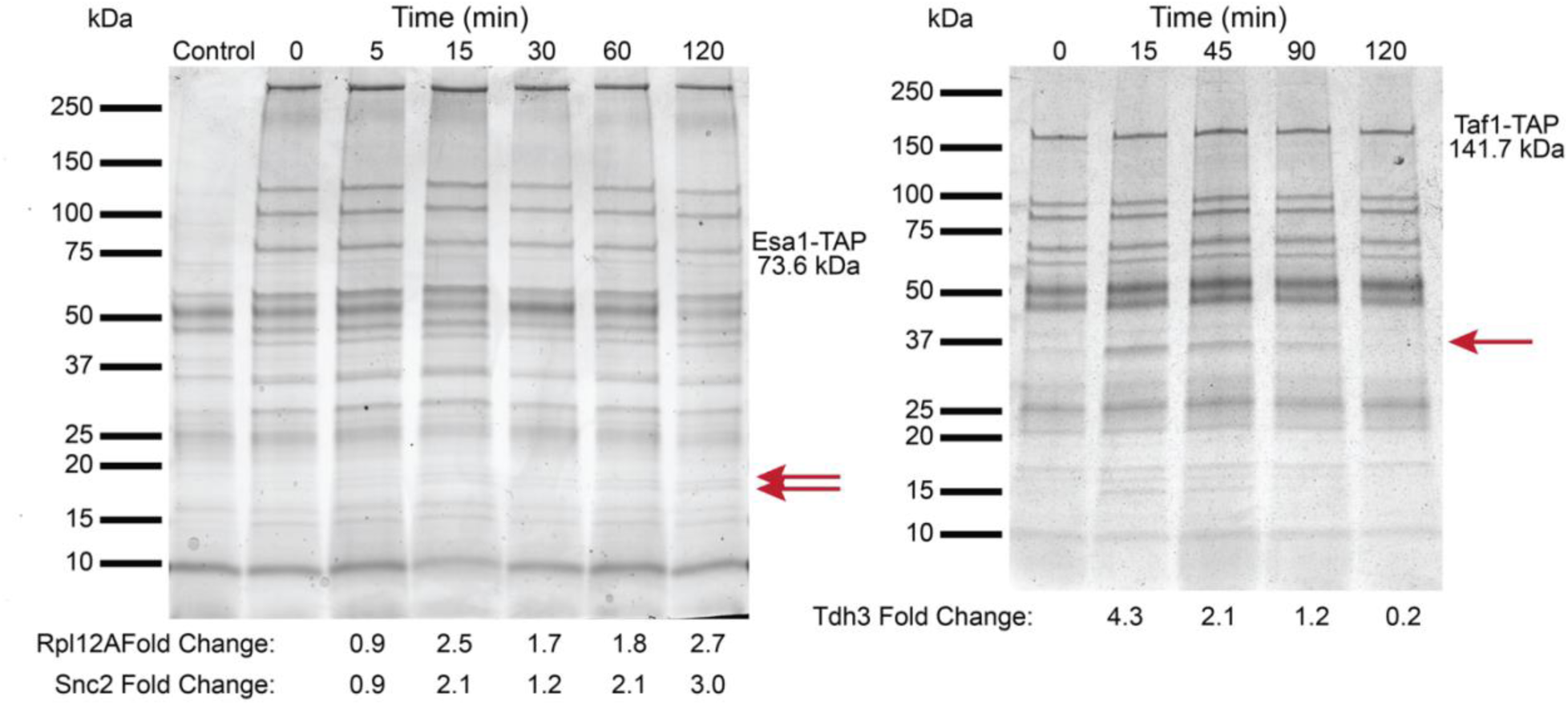
Co-Immunoprecipitation of Esa1-TAP and Taf1-TAP. Co-immunoprecipitation of Esa1 (left) and Taf1 (right) was performed, and proteins were visualized with blue-silver staining. For Esa1 co-IP, two proteins changed in abundance (red arrows) during 25°C to 37°C heat shock after normalization to Esa1-TAP abundance, and were identified via MS as the ribosomal protein Rpl12A and vesicle membrane receptor protein Snc2. For Taf1 co-IP, one protein changed in abundance during heat shock and was identified as the glyceraldehyde-3-phosphate dehydrogenase isoform Tdh3.

Finally, we investigated whether heat shock could induce changes in KAT or KDAC localization that could affect their access to different substrate pools. Using GFP fusion strains, we systematically performed fluorescence microscopy on each of the potential regulatory KATs and KDACs identified via our clustering analysis. Among all enzymes examined, only KDAC She1 showed localization changes, moving from the nucleus to the cytoplasm after 60 minutes at 37°C (Figure 9 and Figure S5). This timing correlates well with the deacetylation of many cytoplasmic proteins in our dataset, particularly those involved in protein folding. Previous work has shown that She1 can deacetylate the cytoplasmic chaperone Hsp90 [44], suggesting its relocalization could be important for deacetylating certain stress-responsive proteins during heat shock.

**Figure 9.**
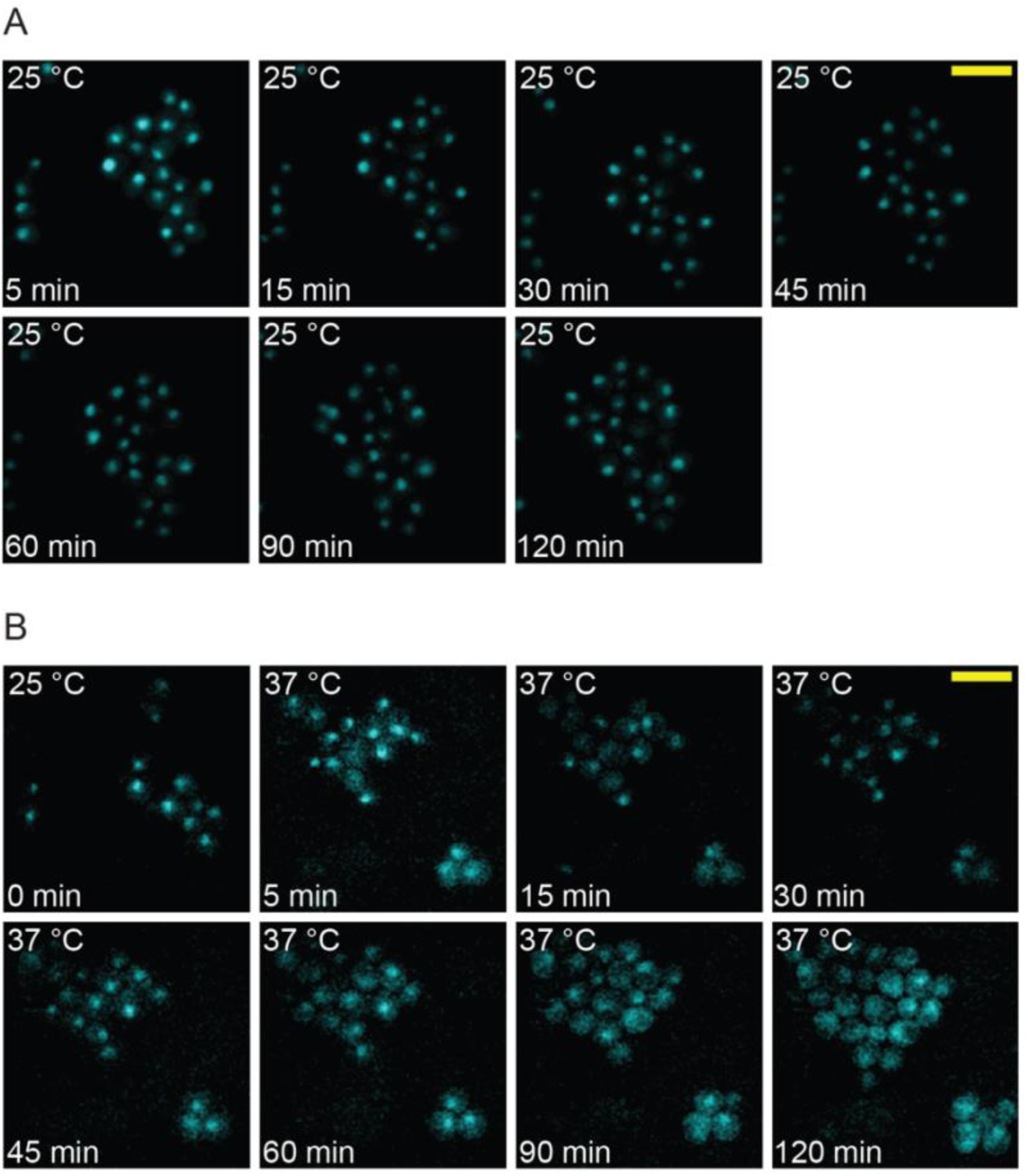
Hda1 relocalizes to the cytoplasm after 45 minutes of heat shock at 37°C. Strains carrying a GFP-tagged Hda1 were visualized for 2 hours at A) 25°C (control treatment) or at 37°C (heat shock) to examine Hda1-GFP localization.

Altogether, our findings suggest that cells likely employ multiple, complementary mechanisms to regulate protein acetylation during stress. While global metabolite changes likely occur too early to drive the bulk of acetylation dynamics, changes in the protein levels of specific KATs and KDACs may play a role, and we additionally have some potential examples of heat shock potentially affecting KAT protein-protein interactions (Esa1 and Taf1) and KDAC relocalization (She1). We examined the growth of main potentially implicated non-essential KAT (*eaf1*Δ and *hat1*Δ) and KDAC (*sir2*Δ and *she1*Δ) mutants, plus the potential interaction partners from co-immunoprecipitation experiments (*rpl12a*Δ, *snc2Δ,* and *tdh3Δ*) (Figure S6). The *eaf1*Δ mutant (which unlike Esa1 is a non-essential component of the NuA4 acetyltransferase complex) had a strong defect during growth at 40°C, while the suspected dynamic interaction partner mutant (*rpl12aΔ*) also had a strong defect. The *tdh3Δ* mutant (encoding the suspected dynamic interaction partner for Taf1) had a small fitness defect at 40°C. In contrasts, the *she1Δ* mutant had a slight but reproducible fitness advantage at 40°C. Additionally, *gcn5Δ* [4, 87–89] and *elp3*Δ [89–95] KAT mutants have reported heat sensitivity phenotypes across multiple studies, as does the *rpd3Δ* KDAC mutant [4, 96]. These phenotypic results are consistent with observed dynamic acetylome remodeling having a functional impact on cellular fitness during heat stress. Ultimately, the regulation of acetylome remodeling is likely complex, and future acetyl-proteomic studies on KAT and KDAC mutants during heat shock would help better define their specific roles.

### Synthesis and Implications

In this study, we used acetyl-proteomics to examine the dynamics of acetylome remodeling during the yeast heat shock response. We provide genetic evidence that protein acetylation impacts the activity of the key co-chaperone Sti1, and we suggest some potential mechanisms for how acetylome remodeling during stress may be regulated. Our findings have potentially broad implications for understanding how acetylation may regulate cellular stress responses. First, the timing of acetylation changes—peaking near 90 minutes—suggests that this modification provides an additional layer of regulation that complements rather than precedes gene expression responses. This differs markedly from phosphorylation changes, which occur within minutes of heat shock [67], indicating that acetylation likely serves a distinct regulatory role during stress, perhaps fine tuning the activities of already translated proteins.

One caveat is that very early acetylation events may be missed in our study, as our first time point was 5 minutes. For example, increased histone acetylation at the yeast *HSP82* promoter has been observed within the first minute of heat shock [97]. While this does not affect our interpretation of bulk dynamics, it is possible that a subset of proteins with dynamic acetylation during heat shock show remodeling before our first measured time point. Our findings also inform upon ongoing debates about the functionality of protein acetylation. Some have suggested that many acetylation events represent ‘noise’ rather than regulated signaling [98, 99], supported by the idea that many non-nuclear acetylation sites are of low stoichiometry [41]. Others have argued that low acetylation stoichiometry should not be dismissed out of hand as unimportant [100], as localization of protein subpopulations can still be important for regulation. Future studies examining the stoichiometry of dynamic acetylation events during heat shock are clearly warranted, as sites with high stoichiometry either before or after stress should be enriched for functional importance. With that caveat in mind, this study suggests that some of the observed dynamic acetylation events during heat shock likely play a bona fide regulatory role. The strong enrichment of stress defense functions for proteins with multiple changing acetyl lysine residues, combined with the presence of both increasing and decreasing acetylation on individual proteins, suggests coordinated regulation rather than noise. Indeed, the presence of both increasing and decreasing acetylation marks on individual proteins suggests sophisticated control mechanisms reminiscent of the histone and chaperone codes, with combinatorial control of activating and repressive acetyl marks likely being a key feature of the acetylome. We do note that one major caveat of acetyl-proteomics is that some peptides cannot be unambiguously mapped to a single protein, which most commonly happens with paralogs (Table S3). A subset of proteins had ambiguously assigned opposing acetyl-marks, which could reflect paralog-specific differences (e.g., one paralog becoming fully acetylated while another is deacetylated), potentially serving to coordinately activate and repress specific paralogs rather than reflect combinatorial regulation within a single protein. However, for paralogs with very high levels of sequence identity, it seems unlikely that regulatory KATs and KDACs could distinguish between paralogs. Nonetheless, targeted mass spectrometry and biochemical dissection of individual proteins are needed to distinguish between these possibilities. We do observe other examples of proteins with unambiguous dynamic acetylation events that are temporally separated in a way that could suggest coordinated regulation. For example, Hsp60 displays dynamic deacetylation of one lysine residue that precedes increased acetylation at multiple separate sites (Figure 5), raising the possibility that deacetylation at one site is required for acetylation at others. In other cases, acetylation dynamics of two lysines on the same protein appear as ‘mirror images’ which could suggest mutually exclusive acetyl sites (e.g., Tef1, Rpl5, and Rps6A). We argue that the strong overlap between proteins with changing acetylation during heat shock and those whose gene expression is activated by heat shock suggests that acetylome remodeling during heat shock may help reinforce transcriptional responses, for example, by activating stress-induced proteins and inactivating repressed ones. Future genetic and biochemical studies are clearly warranted to continue to tease apart the functional role of acetylation at more protein targets.

Comparing our dataset with previous acetylome studies reveals both conserved and unique features. Like acetylation changes observed in bacteria [50] and marine invertebrates [51, 52] during stress, we find extensive modification of proteins involved in energy metabolism and protein folding. However, our analysis revealed regulation of additional processes, particularly translation and ribosome biogenesis, that may reflect differences in timing or biology of the study systems. Our identification of proteins showing both increasing and decreasing acetylation on different residues is also notable, as this pattern has not been reported in other systems.

The functional significance of these acetylation changes likely extends beyond their role in stress defense. Many of the proteins showing altered acetylation are involved in core cellular processes like translation and metabolism. Understanding how acetylation affects their activity could provide insight into how cells generally coordinate protein function through PTMs. Overall, this study suggests that acetylome remodeling is a key feature of the yeast heat shock response and likely stress responses more generally. The breadth of proteins affected suggest that acetylation plays a broader role in stress defense than previously appreciated. Further studies examining both mechanism and function will help explain how cells use acetylation to coordinate adaptation to environmental change.

## Materials and Methods

### Yeast Strains

Strains used in this study are listed in Table S1, and plasmids and oligonucleotides are listed in Table S2. All proteomic, metabolite, immunoprecipitation, and GFP localization experiments were performed using BY4741 (S288c background; MATa *his3Δ1 leu2Δ0 met15Δ0 ura3Δ0*) or its derivatives. Immunoprecipitations were performed using the TAP Tagged ORF Collection (MATa *his*3Δ1 *leu*2Δ0 *met*15Δ0 *ura*3Δ0 ORF::TAP-HIS3) (GE Dharmacon). Protein localization was visualized using GFP fusions obtained from the Yeast GFP Clone Collection (MATa *his*3Δ1 *leu*2Δ0 *met*15Δ0 *ura*3Δ0 ORF::GFP-hisMX6) (Invitrogen). Homozygous gene deletion mutant experiments were performed using the BY4743 background (S288c; *MATa/α his3Δ1/his3Δ1 leu2Δ0/leu2Δ0 lys2Δ0/LYS2 met15Δ0/MET15 ura3Δ0/ura3Δ0*) with the exception of *sir2Δ* which was performed in the BY4741 background. Deletion mutants were obtained from the Yeast Knockout (YKO) Collection (Open Biosystems, now Horizon Discovery). Deletions were verified with diagnostic PCR that included primer pairs that anneal to the MX cassette that replaces the ORF and a sequence upstream of the deletion (verifying the ‘scar’), and primer pairs that anneal within the deletion (verifying gene absence). To generate mimetic mutations in *STI1,* the gene was cloned into pAG36 [101], and site directed mutagenesis was then performed using the QuikChange II Site-directed mutagenesis kit (Agilent). Plasmid mutations were verified via Sanger sequencing performed by Eurofins Genomics.

### Acquired Thermotolerance and Growth Assays

Acquired stress resistance assays were performed as described in [102, 103] with the modification of using a thermal cycler (Bio-Rad DNA Engine) with a gradient function to simultaneously assay survival at multiple temperatures. Briefly, cells were grown with orbital shaking (270 rpm) at 25°C to saturation in YPD (1% yeast extract, 2% peptone, 2% dextrose), and then sub-cultured and grown for at least 8 generations at 25°C to mid-exponential phase (OD_600_ of 0.3 – 0.6 on a Unico spectrophotometer). Cells were then split into two samples and collected by mild centrifugation at 1,500 x *g* for 3 min. The ‘mock’ pretreatment samples were resuspended in prewarmed 25°C media and then incubated at 25°C for 1 h with orbital shaking (270 rpm), while the mild heat shock (‘primary’) pretreatment samples were resuspended in prewarmed 37°C media and then incubated at 37°C for 1 h with 270 rpm orbital shaking. When using cycloheximide (10 µg/ml) or the KDAC inhibitors Trichostatin A (10 µM) and nicotinamide (5 mM), they were added to samples 20 min prior to pretreatment and then throughout the 1-hour pretreatment. Mock and primary-stress pretreated cells were then collected by centrifugation at 1,500 x *g* for 3 min, resuspended in fresh media with no inhibitors to an OD_600_ of 0.6, and transferred to a sterile PCR plate. The plate was sealed with breathable Rayon film (VWR) and cells were incubated in thermal cycler under a gradient of 42°C to 48°C for 1 h. Cells were then serially diluted, plated onto YPD, and incubated at 30°C, and colony-forming units (CFUs) were used to quantify viability. Percent viability was calculated relative to the 42°C samples within each treatment condition.

For growth curve analysis of deletion strains and mimetic mutants, ∼4 transformants were grown overnight with orbital shaking (270 rpm) at 30°C to saturation on synthetic complete (SC, [104]) media, or on SC lacking uracil (SC -Ura) for strains carrying plasmids. Cultures were diluted 1:500 into SC -Ura in 24-well plates, and growth was monitored by measuring the OD_600_ every 15 min in an Eon microplate spectrophotometer (BioTek) with continuous orbital shaking at 40°C. Cells were grown 24 h for control experiments (30°C) or 48 h for heat stress experiments (40°C). For the mimetic experiments, 6 biological replicates were performed. For all other growth experiments, biological triplicates were performed.

### Yeast Sample Preparation for Acetyl-Proteomics

Global acetylation dynamics were obtained by subjecting cells to a 25°C to 37°C heat shock across a 240-min time course as described in [105]. Briefly, two 1 L cultures of exponentially growing cells (OD_600_ of 0.3 – 0.6) were pooled and then split across 3 flasks. Two flasks received 250 ml of culture, while the third flask received 250 ml of the culture and 250 ml of 25°C YPD (this dilution was necessary for cells to maintain exponential growth during the 90-, 120-, and 240-min time points). Heat shock was initiated by adding an equal volume of 55°C YPD to the cells, which immediately brought the final temperature to 37°C. An unstressed control sample was collected, and heat-shocked cells were incubated at 37°C until collection at 5, 10, 15, 30, 45, 60, 90, 120, and 240-min post-heat shock. Cells were collected onto cellulose nitrate filters by vacuum filtration of 120 ml of culture. Cells were immediately scraped from the filters into liquid nitrogen and stored at -80°C until processing. Three independent heat shock time course collections were performed, resulting in biological triplicates.

### TMT-10-Plex Sample Preparation

For each sample, 90% was used for acetyl-enrichment, while 10% was reserved for total proteome analysis for normalizing changes in acetyl-peptide abundance to the changes in abundance of total proteins. Our processing and analysis of the unenriched samples was previously described in [105]. Briefly, samples were resuspended in fresh 6M guanidine hydrochloride and 100 mM Tris pH 8.0. Rapid lysis at high temperature was then performed (100°C for 5 min, 25°C for 5 min, and 100°C for 5 min) to stabilize PTMs in the absence of specific inhibitors [106]. Samples were precipitated by adding nine volumes of 100% methanol, vortexing, and centrifuging at 9000 x *g* for 5 min. The protein pellets were air dried for 5 min and then resuspended in 8 M urea.

Protein samples (∼5 mg total) were diluted to 2 M urea 100 mM Tris pH 8.0, and digested with a 1:50 ratio of trypsin overnight at 25°C (to prevent carbamylation of free amines [107, 108]), with gentle mixing in the presence of 2.5 mM tris(2-carboxyethyl)phosphine (TCEP) and 10 mM chloroacetamide for reduction and alkylation. Digestion was quenched with 0.6% trifluoroacetic acid (TFA) to a pH less than 2, and peptides were desalted with Sep Pak C18 columns and lyophilized. A detailed protocol for peptide desalting can be found on the protocols.io repository under DOI dx.doi.org/10.17504/protocols.io.3hegj3e. Peptides were resuspended in 200 mM tetraethylammonium bromide (TEAB) to a final concentration of ∼8 μg, quantified with Pierce Colorimetric Peptide Assay, and diluted to 5 μg/ μl in 200 mM TEAB. Five hundred µg of each sample was labeled with one of ten tandem mass tag (TMT) isobaric labels (Thermo Fisher Scientific). Each sample was mixed with a separate TMT label reconstituted in 50 µl acetonitrile and incubated at room temperature for 1 h. Labeling was quenched with 8 µl 5% hydroxylamine for 15 min, and samples were combined, desalted, and lyophilized. Labeling efficiency was monitored by performing Mascot searches with the TMT-10 modification mass as a variable modification instead of fixed. As reported in [105], the labeling efficiencies were 96.3%, 97.4%, and 97.5% for biological replicates 1, 2, and 3, respectively. A detailed protocol for cell lysis and TMT labeling can be found at dx.doi.org/10.17504/protocols.io.3g9gjz6.

### Basic pH Fractionation and Acetyl-Lysine Enrichment

The processing of the ‘control’ samples used for quantifying changes in total protein abundance during heat shock was described in [105]. For acetyl-lysine enrichment samples, 450 µg of pooled peptides were fractionated for each biological replicate. Peptides were dissolved in 10mM ammonium bicarbonate (pH 10), and fractionated using an Agilent 300 Extend C18 column (770995-902) in a gradient of 0-100% acetonitrile in 10mM ammonium bicarbonate (pH 10) on a Shimadzu LC-20AP system. The first 10 min (flow-through) were collected as fraction 1 and the remaining fractions were collected for 2.5 minutes for a total of 10 fractions, and then a final fraction was collected for 2.5 minutes and added to fraction 1. The 10 fractions were frozen in liquid N_2_ and lyophilized prior to acetyl-lysine enrichment.

Each fraction was dissolved in 1.4 ml IAP buffer (50 mM MOPS pH 7.2, 10 mM Na_2_HPO_4_, 50 mM NaCl), transferred to a tube containing 40 µl of PTMScan Acetyl-Lysine Motif [Ac-K] IAP Beads (Cell Signaling Technology, 13362S), and incubated at 4°C with gentle rotation for 2 h. Samples were washed with IAP buffer and water and eluted with 0.15% TFA according to the manufacturer’s protocol. Eluted samples were stored at -80°C prior to mass spectrometry.

### LC-MS/MS Data Analysis

Pooled acetyl-enriched peptides were then fractionated again on a 100 mm x 1.0 mm Acquity BEH C18 column (Waters) using an UltiMate 3000 UHPLC system (Thermo Fisher Scientific) with a 40 min gradient from 99:1 to 60:40 buffer A:B ratio under basic (pH 10) conditions (buffer A = 0.05% acetonitrile, 10 mM NH_4_OH; buffer B = ∼100% acetonitrile, 10 mM NH_4_OH (22% aqueous NH_4_OH diluted into 100% acetonitrile). The 96 individual fractions were then consolidated into 24 superfractions using the concatenation scheme described in [109] (i.e., 1 + 25 +49 +73, 2 + 26 +50 +74, etc.).

Superfractions were loaded on a 150 mm x 0.075 mm column packed with Waters C18 CSH resin. Peptides were eluted using a 70-min gradient from 96:4 to 75:25 buffer A:B ratio into an Orbitrap Fusion Lumos mass spectrometer (Thermo Fisher Scientific). MS acquisition consisted of a full scan at 120,000 resolution, a maximum injection time of 50 ms, and an AGC target of 7.5 x 10^5^. Selection filters consisted of monoisotopic peak determination, charge state 2-7, intensity threshold of 2.0 x 10^4^, and mass range of 400-1200 *m*/*z*. The dynamic exclusion length was set to 15 s. The data-dependent cycle time was set for 2.5 s. Isolation widths were 0.7 Da for MS^2^ and 2 Da for the MS^3^ scans. Selected precursors were fragmented using CID 35% with an AGC target of 5.0 x 10^3^ and a maximum injection time of 50 ms. MS^2^ scans were followed by synchronous precursor selection of the 10 most abundant fragment ions, which were fragmented with HCD 65% and scanned in the Orbitrap at 50,000 resolution, an AGC target of 5.0 x 10^4^, and a maximum injection time of 86 ms.

Proteins were identified by database search using MaxQuant50 (Max Planck Institute) using the Uniprot *S. cerevisiae* database from October 2014 [110], with a parent ion tolerance of 3 ppm and a fragment ion tolerance of 0.5 Da. Carbamidomethylation of cysteine residues and TMT-labelling (+229.16) of peptide N-termini were used as fixed modifications. Acetylation of lysine residues and protein N-termini, and oxidation of methionine were selected as variable modifications. Mascot searches were performed using the same parameters as above, and were imported into Scaffold software (v4) [111]] and filtered for protein and peptide false discovery rates of 1% across all three biological replicates. Ambiguous acetyl-peptides mapping to more than one protein were assigned to all matched proteins, and all identified acetyl-lysines can be found in Table S3. Data normalization and analyses were performed using R and scripts are provided in [105] and under Zenodo doi: 10.5281/zenodo.18462377. Spectra containing missing values in any channel were excluded from quantitation. Spectra were further filtered to include only high-scoring peptide-spectrum matches (Mascot ion score cutoff of >15). Reporter ion intensities were log_2_ transformed, mean-centered for each spectrum, then median-centered for each channel to control for mixing. Peptide and protein quantitative values were obtained by taking the mean of the quantitative values for all spectra mapping to the peptide or protein. All raw mass spectrometry data and MaxQuant search results have been deposited to the ProteomeXchange Consortium [112] via the PRIDE [113] partner repository with the dataset identifier PXD014552 and 10.6019/PXD014552 for the total proteome data [105], and PXD063569 and 10.6019/PXD063569 for the acetyl-enrichment data reported here.

### Acetyl-Proteomic Statistical Analyses

To identify changes in acetyl-lysine peptides and total protein abundance during heat shock, unstressed cells were used as a reference within each TMT-10 replicate to calculate relative log_2_ abundance changes, which is a strategy that has been used successfully to identify changes in relative abundance across multiple TMT experiments [105, 114]. To identify high-confidence changes in acetyl-lysine peptide abundance, only acetyl-lysine peptides and total proteins that were identified in all time points across all three biological replicates were included. To control for changes in acetylation abundance that only reflect underlying changes in protein abundance over the time course, acetyl-peptide log_2_ fold changes were further normalized to changes in total protein abundance at each time point. Significant changes in acetyl-lysine peptide and total protein abundance were identified by performing an empirical Bayes moderated *t-*test using the BioConductor package Limma v 3.36.2 [115] and Benjamini-Hochberg FDR correction [116], with FDR <0.05 being used as a significance cutoff.

Hierarchical clustering was performed using Cluster 3.0 (http://bonsai.hgc.jp/∼mdehoon/software/cluster/software.htm), with Euclidian distance as the metric with centroid linkage [117], with experimental weights using a cutoff value of 0.4 and an exponent value of 1. Heat maps were visualized using Java TreeView v. 1.1.6r4 [118], and clusters were manually collapsed using a minimum Pearson correlation of 0.9.

Gene ontology (GO) functional enrichments were performed using the Princeton GO-TermFinder [119] with Bonferroni-corrected *P*-values < 0.01 taken as significant. KAT and KDAC enrichments were performed by extracting known protein-protein interactions from the *Saccharomyces* Genome Database [120], and applying Fisher’s exact test with *P*-values < 0.05 taken as significant. Motif analysis was performed using the motif-x algorithm in the modification motifs (MoMo) tool [121] found in the MEME suite [122]. When comparing acetylation dynamics with mRNA levels during heat shock, microarray time course data was obtained from [123] and RNA-seq data during the peak of the transcriptional response was obtained from [69].

### Metabolite Extraction and Analyses

To measure acetyl-CoA, free CoA, and nicotinamide levels during heat shock, cells were grown to mid-exponential phase (OD_600_ of 0.4-0.6) for at least 8 generations in YPD at 25°C with 200 rpm orbital shaking, and then samples were aliquoted into fresh prewarmed 55°C to immediately bring the temperature to 37°C. Heat-shocked cells were then incubated at 37°C with 200 rpm orbital shaking until collection. An unstressed sample was collected as well as heat-shocked samples at 5, 15, 30, 60, and 120 min. For all samples, 1 x 10^8^ cells were collected via vacuum filtration as described above for the acetyl-proteomic sample collections.

Metabolite extraction was performed as described in [124]. Briefly, filters containing cells were placed onto -20°C-chilled extraction solvent (40% acetonitrile, 40% methanol, and 20% water), and metabolites were extracted at -20°C for 15 minutes. The filter was washed 10 times with the pooled solvent, and solvent and cellular debris were collected and centrifuged at 21,130 x *g* for 5 min at 4°C, and the supernatant was transferred and stored at -80°C prior to MS.

Quantitation of nicotinamide, acetyl-CoA), and free-CoA) was performed via LC/MS/MS using a multiple reaction monitoring (MRM) approach. Samples were separated at 25°C by HPLC using a 2.1 mm x 150 mm Waters Atlantis T3 column and an Agilent 1100 Binary Pump, using a 30-min gradient: 100:1 buffer A:B ratio for 3 min, 100:0 to 30:70 A:B over 17 min, 30:70 A:B for 3 min, 100:0 buffer A:B over 1 min, 100:0 buffer A:B for 6 min (buffer A = 25 mM ammonium formate, pH 7.5; buffer B = 100% acetonitrile). MS/MS detection was performed on a Thermo TSQ Quantum Discovery Max triple quadrupole instrument with positive ion electrospray ionization. The ESI source was a HESI-II probe with vaporizer temperature of 300°C and ionization voltage of 4200V. The sheath gas was set to 35 (arbitrary units), with an ion sweep gas of 5 and auxiliary gas of 10. The inlet capillary was held at 300°C with a capillary offset of 35V.

Pure standards of acetyl-CoA, CoA, and nicotinamide were obtained from Sigma Aldrich. Standards were used to generate ‘spike ins’ for normalization of each sample at 0 µM (water alone), 2 µM, and 20 µM. Data analysis was performed using Quantitative Analysis software (Thermo Fisher) to integrate peak areas for each analyte. Peak areas consisted of the summed signal for all MRM transitions for each analyte. Calibration curves were created from the spike in samples and were then used to calculate analyte concentrations for each sample.

To measure NAD:NADH ratios, cells were processed as described for metabolite LC-MS-MS, with the exception of lysis being performed by resuspending cells in lysis buffer (50% PBS and 50% 0.2 N NaOH, 3x protease inhibitor cocktail (VWR A32963)), freezing in liquid N_2_, and grinding with a mortar and pestle. Lysed cells were thawed and centrifuged at 21,130 x *g* for 5 min at 4°C to pellet debris. Protein concentration was determined via Bradford assay, and samples were diluted to 10 μg / ml in lysis buffer. Samples were split to measure NAD^+^ and NADH individually using the NAD/NADH-Glo™ Assay (Promega), according to the manufacturer’s instructions, with luminescence measured using a Synergy H1 microplate reader (Biotek).

### TAP Purification and Fluorescence Microscopy

For TAP purification, cells were collected as described for acetyl-proteomics, with an unstressed 25°C control sample being collected at 15, 45, 90, and 120 minutes. TAP purifications were performed as described in [125]. Briefly, cells were thawed on ice in lysis buffer (20 mM HEPES, pH 7.4, 0.1% Tween 20, 2 mM MgCl_2_, 300 mM NaCl, protease inhibitor cocktail (VWR A32963)), and then lysed via vortexing with acid-washed glass beads (425–600 μm, Sigma-Aldrich) with 6 cycles of 1 min vortexing followed by 2 min on ice. Lysate was cleared by centrifugation at 14,000 x *g* for 20 min at 4°C, and protein was quantified via Bradford assay. Ten mg of sample was incubated with 25 µl of magnetic Dynabeads (Fisher, 14301) crossed-linked to rabbit immunoglobulin G (IgG) (Sigma, I5006), for 2 h with end-over-end incubation at 4°C. After 5 washes with 1 ml 4°C lysis buffer, Dynabeads were suspended in 25 μl loading buffer (50 mM Tris, pH 6.8, 2% sodium dodecyl sulfate (SDS), 0.1% bromophenol blue, 10% glycerol) and proteins were eluted at 65°C for 10 min. The supernatant was collected, β-mercaptoethanol was added to a final concentration of 200 mM, and the samples were incubated at 95°C for 5 min before resolving on a 4 to 12% polyacrylamide gradient gel (Biorad, 4561093). Proteins were fixed in 50% methanol, 40% ddH2O, 10% acetic acid and visualized by blue-silver staining (0.12% Coomassie Blue G-250, 10% ammonium sulfate, 10% phosphoric acid, 20% methanol) [126]. Mock immunoprecipitation controls were performed on an untagged strain to identify non-specific proteins. Individual bands with changing intensity during the heat shock time course not found in the controls were quantified via densitometry using ImageJ [127] [128], with normalization being applied using the intensity of the relevant TAP-tagged protein in each lane. Bands were then subjected to in-gel trypsin digest and MS identification on an Orbitrap Fusion.

For fluorescence microscopy, cells were grown to mid-exponential phase (OD_600_ of 0.4-0.6) for at least 8 generations at 25°C in SC medium (which has lower background fluorescence compared to YPD). Ten µl of each culture was transferred to a 5 mm x 5 mm agarose pad (3% agarose in SC), and attached to a clean coverslip. A chamber was then constructed by adhering the coverslip to a 35 mm Petri dish (Cell E&G, GBD00002-200) using epoxy. The sample was equilibrated at 25°C in a plate incubator for 10 mins, and heat shock was initiated by placing the sample in a plate incubator at 37°C for 5 mins before being transferred to the microscope. The cells were maintained at 37°C using an onstage incubator and imaged at 5, 15, 30, 45, 60, 90, and 120-min post heat-shock. An unstressed control sample was maintained and imaged at 25°C. Fluorescence microscopy was performed on an Olympus IX-73 inverted microscope with an Olympus TIRF 100× N.A. = 1.49 oil immersion objective, and a 488 nm laser from a multilaser system (iChrome MLE, TOPTICA Photonics, New York) was used to excite GFP. Emissions from the fluorescent proteins were collected by the objective and imaged on an EMCCD camera (Andor, Massachusetts) with an exposure time of 30 ms. The effective pixel size of acquired images was 160 nm. Raw images were processed by first subtracting the background using a rolling-ball algorithm in ImageJ [127],with a radius of 50 pixels. The images were then pseudo-colored cyan for visualization. Raw microscopy images can be found as supplementary materials on Zenodo (doi: 10.5281/zenodo.18462377).

## Supporting information

Table S1: Strains and plasmids used in the study

Table S2: Oligonucleotides used in the study

Table S3: Significant dynamic acetyl lysine peptides

Table S4: Functional Enrichments

## Data Availability

All raw mass spectrometry data and MaxQuant search results have been deposited to the ProteomeXchange Consortium [112] via the PRIDE [113] partner repository with the dataset identifier PXD014552 and 10.6019/PXD014552 for the total proteome data [105], and PXD063569 and 10.6019/PXD063569 for the acetyl-enrichment data reported here. Raw images and analysis scripts are provided in Zenodo (doi: 10.5281/zenodo.18462377).

## Author Contributions

Conceptualization: JAL; Formal Analysis: REH-K, AJS, GAB-W, VRK; Investigation REH-K, TNS, SHE, VRK; Methodology: AJS, SDB, SGM, RDE, AJT; Resources: YW, SDB, SGM, RDE, AJT; Supervision: JAL, YW, WPW; Funding Acquisition: JAL, WPW, AJT. Writing – Original Draft: REH-K, JAL; Writing – Review & Editing: REH-K, AJS, TNS, SHE, GAB-W, VRK, YW, SDB, SGM, RDE, WPW, AJT, JAL

## Acknowledgments

We would like to thank Shannon Servoss and Neda Mahmoudi for the use and assistance with the lyophilizer and Ravi Barabote for use of the Shimadzu HPLC. We thank Mark Powell of Agilent, Suresh Kumar, and Srinivas Jayanthi with HPLC protocol development, and Alex Hebert for helpful conversations on TMT proteomic sample processing. We thank Kristen Baetz and Michael Downey for providing Esa1-GFP and for general advice. This work was supported by grants from the National Science Foundation (IOS-1656602 to J.A.L.), an Arkansas Integrative Metabolic Research Center Pilot Grant (J.A.L.) as part of the AIMRC Center for Biomedical Research Excellence (COBRE) grant (5P20GM139768), the National Institutes of Health (GM081766 and GM145834 to W.P.W., and R01CA282198, P20GM121293, and P20GM103429 to A.J.T.), the Arkansas Biosciences Institute (Arkansas Settlement Proceeds Act of 2000), and by the IDeA National Resource for Quantitative Proteomics and NIH grant R24GM137786 (A.J.T.). R.E.H. was partially supported by the University of Arkansas Cell and Molecular Biology Graduate Program.

## Supplemental Figures

**Figure S1.**
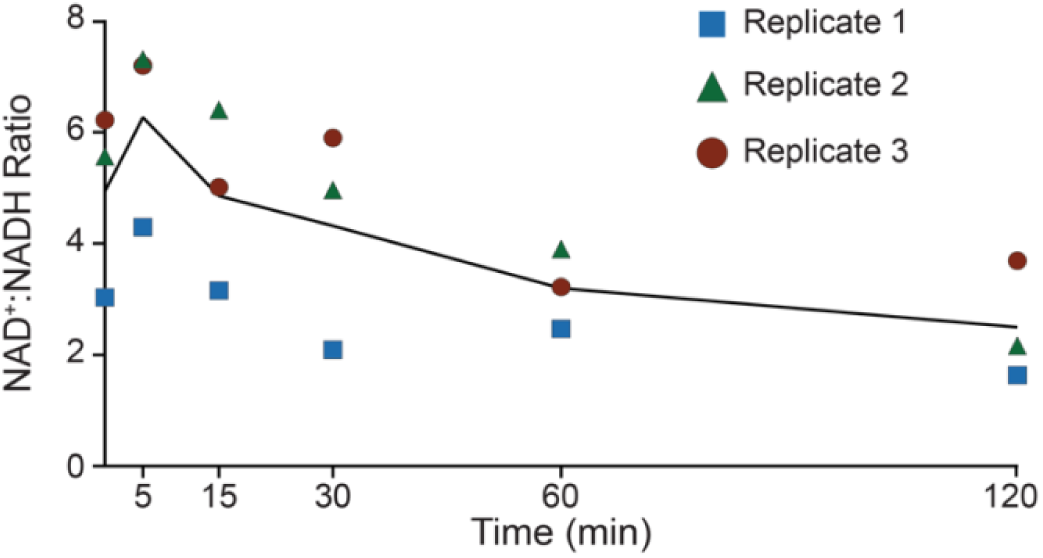
The NAD^+^:NADH ratio initially spikes and then gradually decreases during heat shock. The NAD^+^:NADH ratio was monitored across a 120 min heat shock (25°C to 37°C) via a luminescence assay.

**Figure S2.**
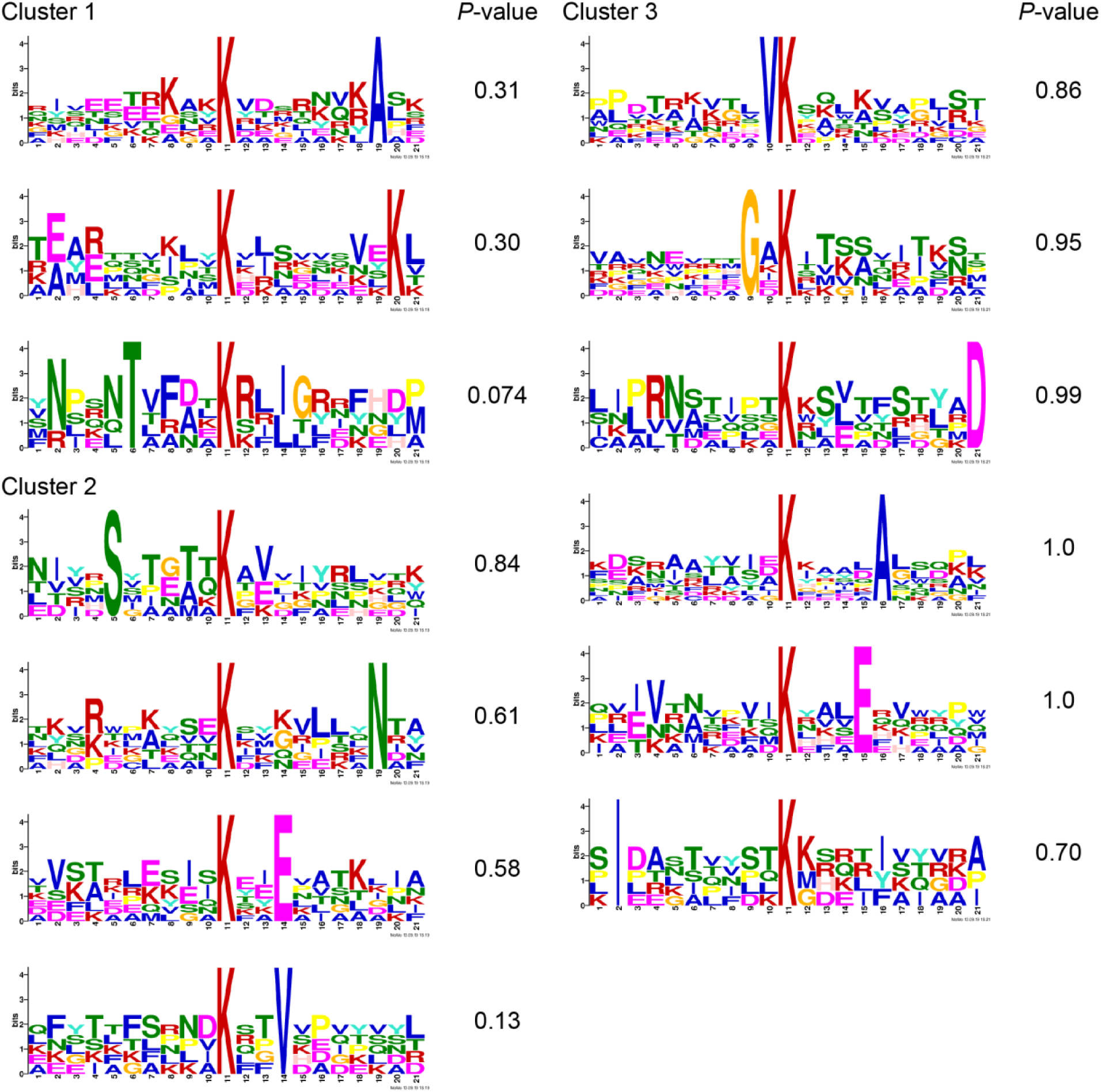
Motif analysis of clusters with increasing acetylation. Motif analysis of Clusters 1-3 from Figure 5A was performed using MoMo.

**Figure S3.**
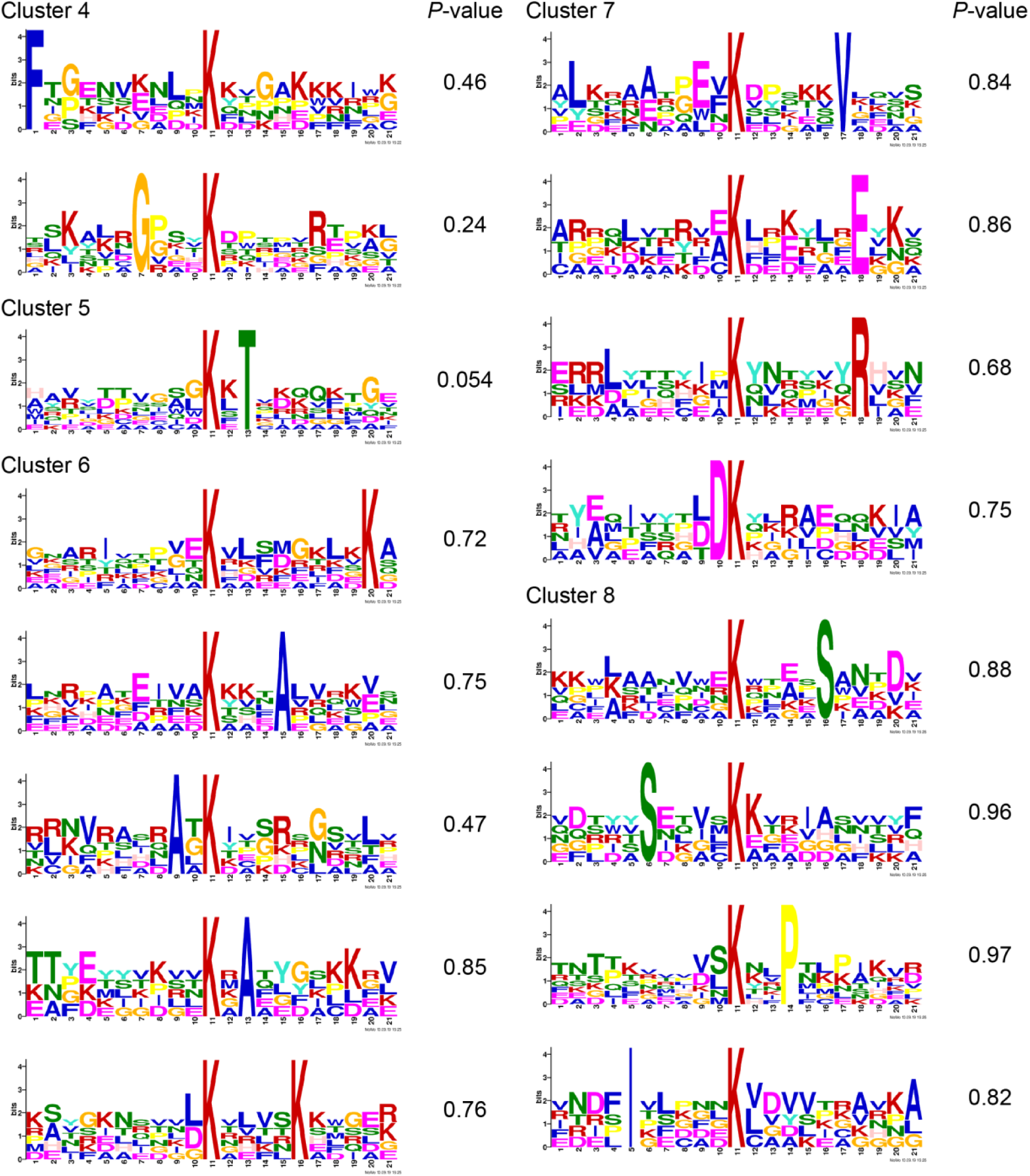
Motif analysis of clusters with decreasing acetylation. Motif analysis of Clusters 4-8 from Figure 5A was performed using MoMo.

**Figure S4.**
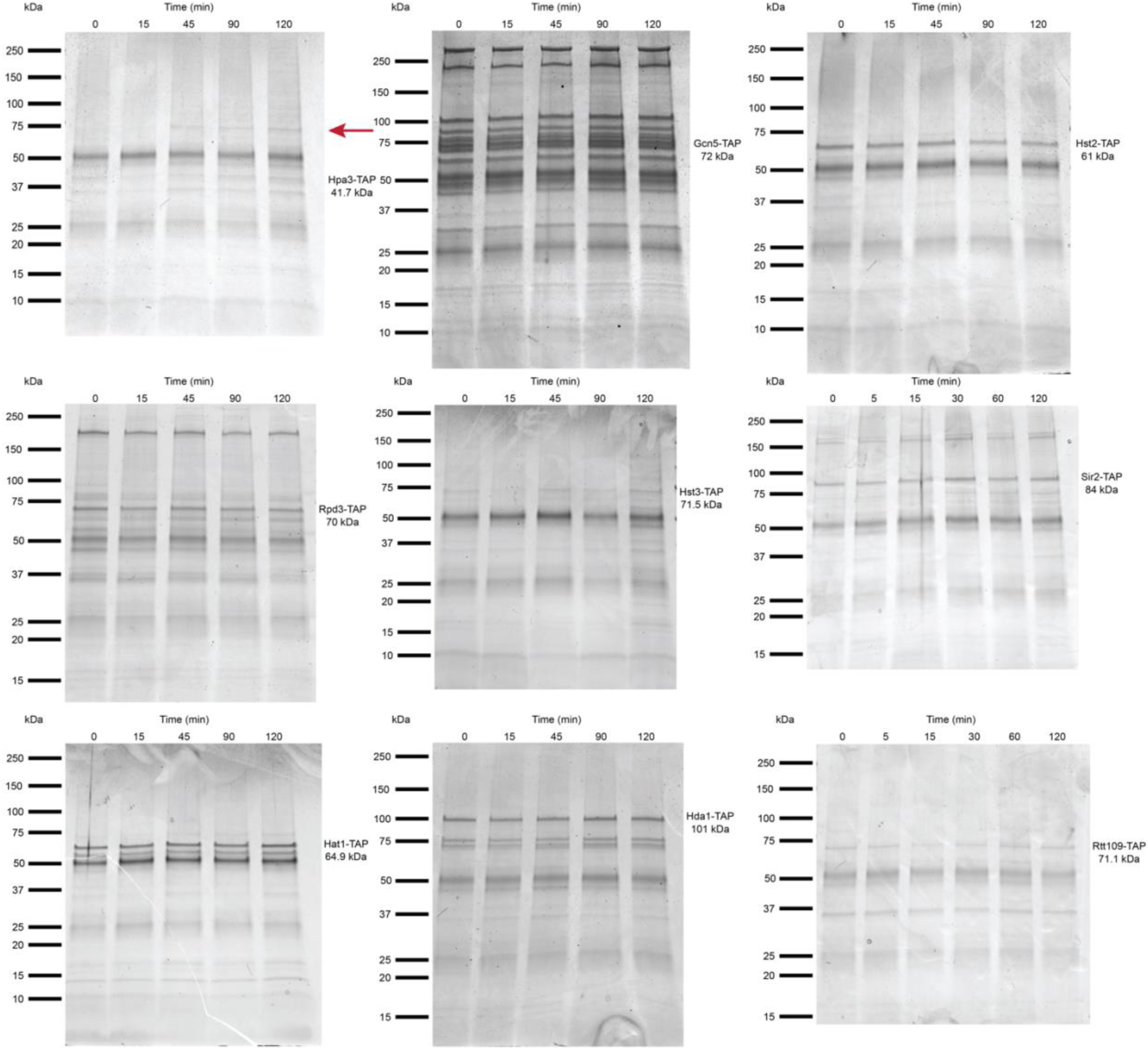
Tap co-immunoprecipitations of KAT and KDAC enzymes and interacting proteins. Identified KAT and KDAC enzymes and interacting proteins were co-immunoprecipitated and visualized with blue-silver staining. One interacting protein changed in abundance for the Hpa3 co-immunoprecipitation (red arrow), but could not be identified. No changes were observed for the other KATs and KDACs shown in the figure.

**Figure S5.**
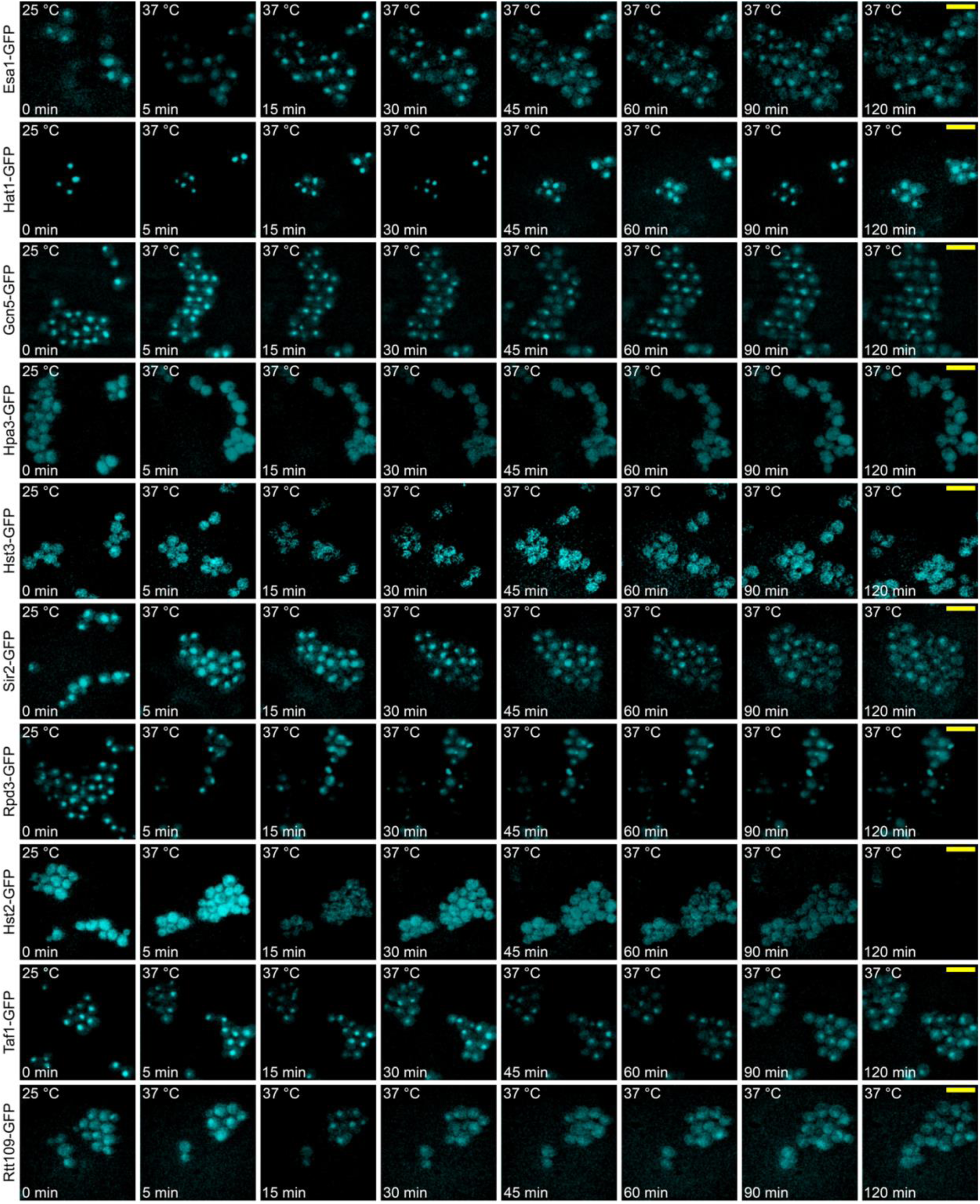
Fluorescence microscopy of KATs and KDACs. Strains carrying the depicted KAT or KDAC-GFP fusions were visualized at 37°C for 2 hours to determine possible changes in localization. None of the depicted proteins show changes in localization.

**Figure S6.**
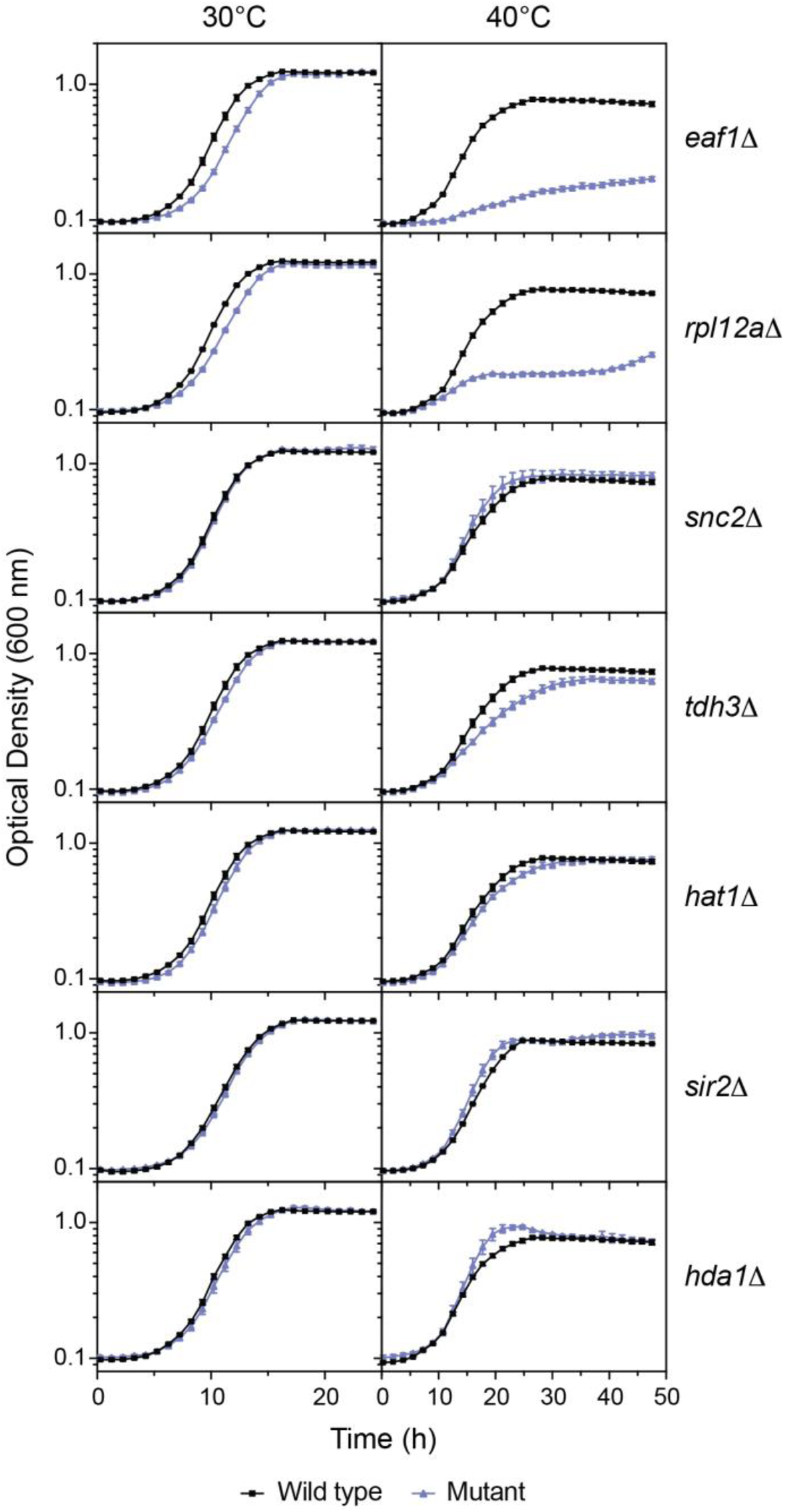
Growth phenotypes of KAT and KDAC mutants or their suspected dynamic interaction partners during heat shock. Growth assays were performed at 40°C comparing wild-type cells with the depicted mutants. The *sir2Δ* mutant was a MATa haploid, and was compared to haploid BY4741. All other mutants were homozygous diploids and were compared to the diploid BY4743. Error bars denote the standard error of 3 biological replicates.

## Supplemental Tables

**Table S1: Strains and plasmids used in the study**

**Table S2: Oligonucleotides used in the study**

**Table S3: Significant dynamic acetyl lysine peptides**

**Table S4: Functional Enrichments**

